# Shared and distinct genetic etiologies for different types of clonal hematopoiesis

**DOI:** 10.1101/2022.03.14.483644

**Authors:** Derek W. Brown, Liam D. Cato, Yajie Zhao, Satish K. Nandakumar, Erik L. Bao, Eugene J. Gardner, Alexander DePaulis, Thomas Rehling, Lei Song, Kai Yu, Stephen J. Chanock, John R. B. Perry, Vijay G. Sankaran, Mitchell J. Machiela

## Abstract

Clonal hematopoiesis (CH) – age-related expansion of mutated hematopoietic clones – can differ in frequency and cellular fitness. Descriptive studies have identified a spectrum of events (coding mutations in driver genes (CHIP), gains/losses and copy-neutral loss of chromosomal segments (mCAs), and loss of sex chromosomes). Co-existence of different CH events raises key questions as to their origin, selection, and impact. Here, we report analyses of sequence and genotype array data in up to 482,378 individuals from UK Biobank, demonstrating shared genetic architecture across different types of CH. These data highlighted evidence for a cellular evolutionary trade-off between different forms of CH, with LOY occurring at lower rates in individuals carrying mutations in established CHIP genes. Furthermore, we observed co-occurrence of CHIP and mCAs with overlap at *TET2, DNMT3A*, and *JAK2*, in which CHIP precedes mCA acquisition. Individuals carrying these overlapping somatic mutations had a large increase in risk of future hematological malignancy (HR=17.31, 95% CI=9.80-30.58, P=8.94×10^−23^), which is significantly elevated compared to individuals with non-overlapping CHIP and autosomal mCAs (P_heterogeneity_=8.83×10^−3^). Finally, we leverage the shared genetic architecture of these CH traits to identify 15 novel loci associated with blood cancer risk.

## Main

Recent studies have reported the frequent occurrence of clonal expansion of post-zygotic mutations in the hematopoietic system, now seen in all human tissues but at different attained frequencies.^1–6^ Initially, clonal expansion was recognized by the presence of skewed X chromosome inactivation.^7,8^ Subsequent studies have revealed the presence of mosaic chromosomal alterations (mCAs), including frequent loss of chromosomes X and Y, in a subset of hematopoietic cells. Most recently, clonal expansion of recurrent somatic driver mutations observed in hematologic malignancies have been identified in individuals with otherwise normal hematologic parameters, a condition known as clonal hematopoiesis of indeterminate potential (CHIP). These somatic alterations can predispose to either myeloid or lymphoid malignancies but do not necessarily progress; in other words, many otherwise healthy individuals are observed to have CHIP.^9^ Moreover, recent studies have shown the additive impact of mCAs and CHIP mutations on predisposition to blood cancers, with respect to overall risk for a primary hematologic malignancy and also in the setting of therapy-associated myeloid malignancies.^10,11^

Large studies have begun to reveal how germline genetic variants can increase risk for acquisition of CH but have yet to systematically investigate the co-existence of events. Moreover, the biologic mechanisms of hematopoiesis that confer risk have not been well-characterized. Here, we utilized a range of genotyping and sequencing data from large-scale studies of genetic susceptibility of types of clonal hematopoiesis (*e*.*g*., mCAs, CHIP), hematologic malignancies *(e*.*g*., myeloproliferative neoplasms (MPNs)), and hematopoietic phenotypes to assess genetic and phenotypic relationships between these distinct but potentially related phenotypes to gain new insights into the underlying mechanisms and consequences of the range of clonal hematopoietic expansions.

## Results

### CH states display shared genetic and phenotypic relationships

We began by investigating the co-existence of different types of CH: loss of chromosome Y (LOY) in men, loss of chromosome X (LOX) in women, autosomal mCAs including gains, losses, and copy neutral loss of heterozygosity (CNLOH), CHIP, and MPNs (**Figure 1**) and subsequently examined associations with germline susceptibility variants, in anticipation of discovery of shared elements. Genome-wide association study (GWAS) summary statistics for each type of CH were analyzed and pairwise genetic correlations between traits were computed (**Online Methods**). Using the high-definition likelihood (HDL) method, positive genetic correlations were observed between LOY and LOX (ρ= 0.23, P= 5.53×10^−9^), LOY and MPN (ρ= 0.35, P= 1.74×10^−4^), autosomal mCAs and MPN (ρ= 0.57, P= 1.83×10^−3^), and CHIP and MPN (ρ= 0.48, P= 4.32×10^−3^) (**Figure 2, Supplemental Table 1**). We repeated genetic correlation analyses using linkage disequilibrium score regression (LDSC) (**Online Methods**) and likewise observed positive genetic correlations for LOY with both LOX (ρ= 0.30, P= 4.09×10^−5^) and MPN (ρ= 0.21, P= 1.67×10^−2^) (**Supplemental Figure 1, Supplemental Table 2**). The genetic correlation between autosomal mCAs and MPN, and CHIP and MPN had the same direction of effect as found with HDL (**Supplemental Figure 1, Supplemental Table 2**).

**Figure 1.**
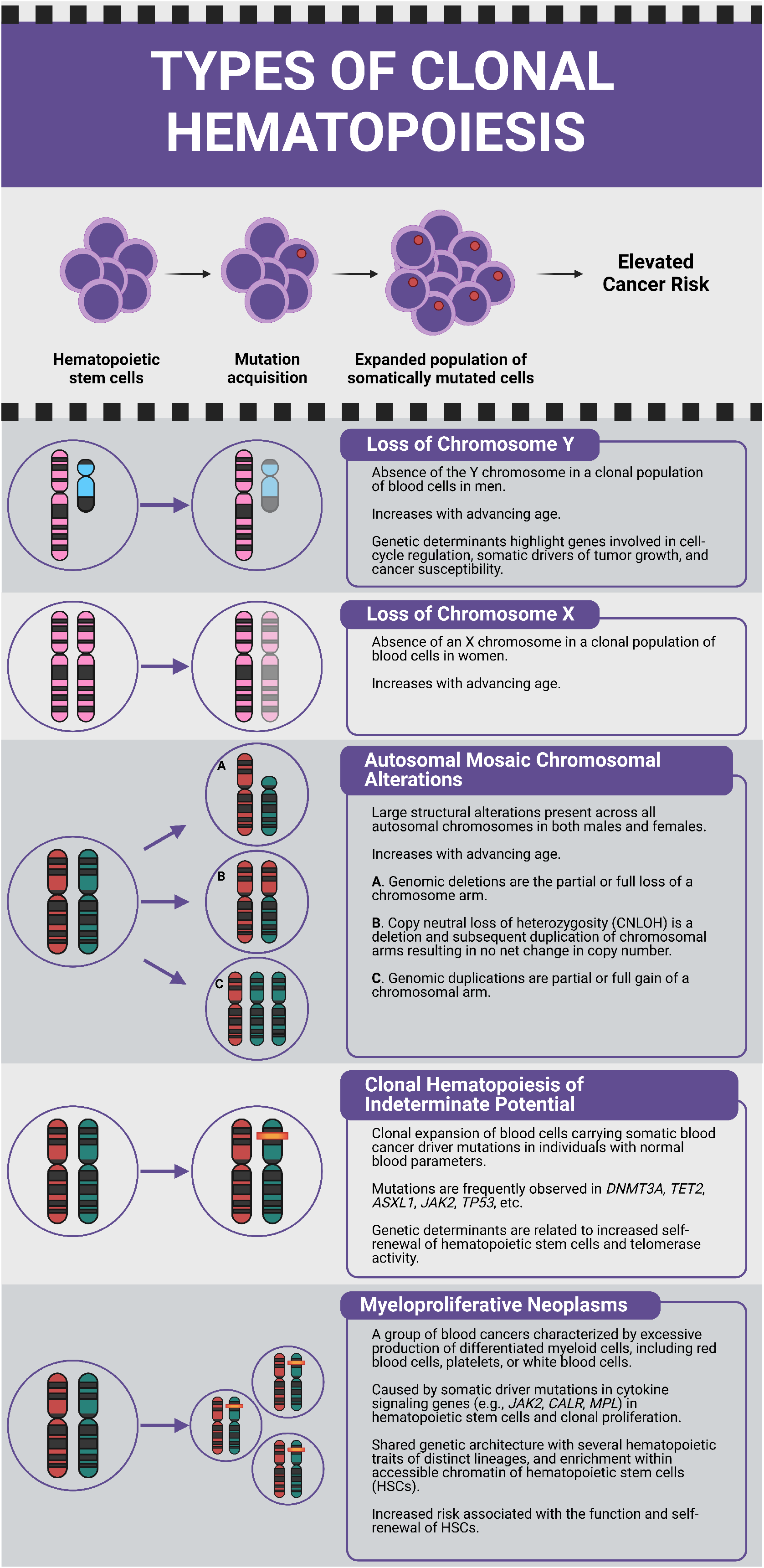
Description of each type of clonal hematopoiesis.

**Figure 2.**
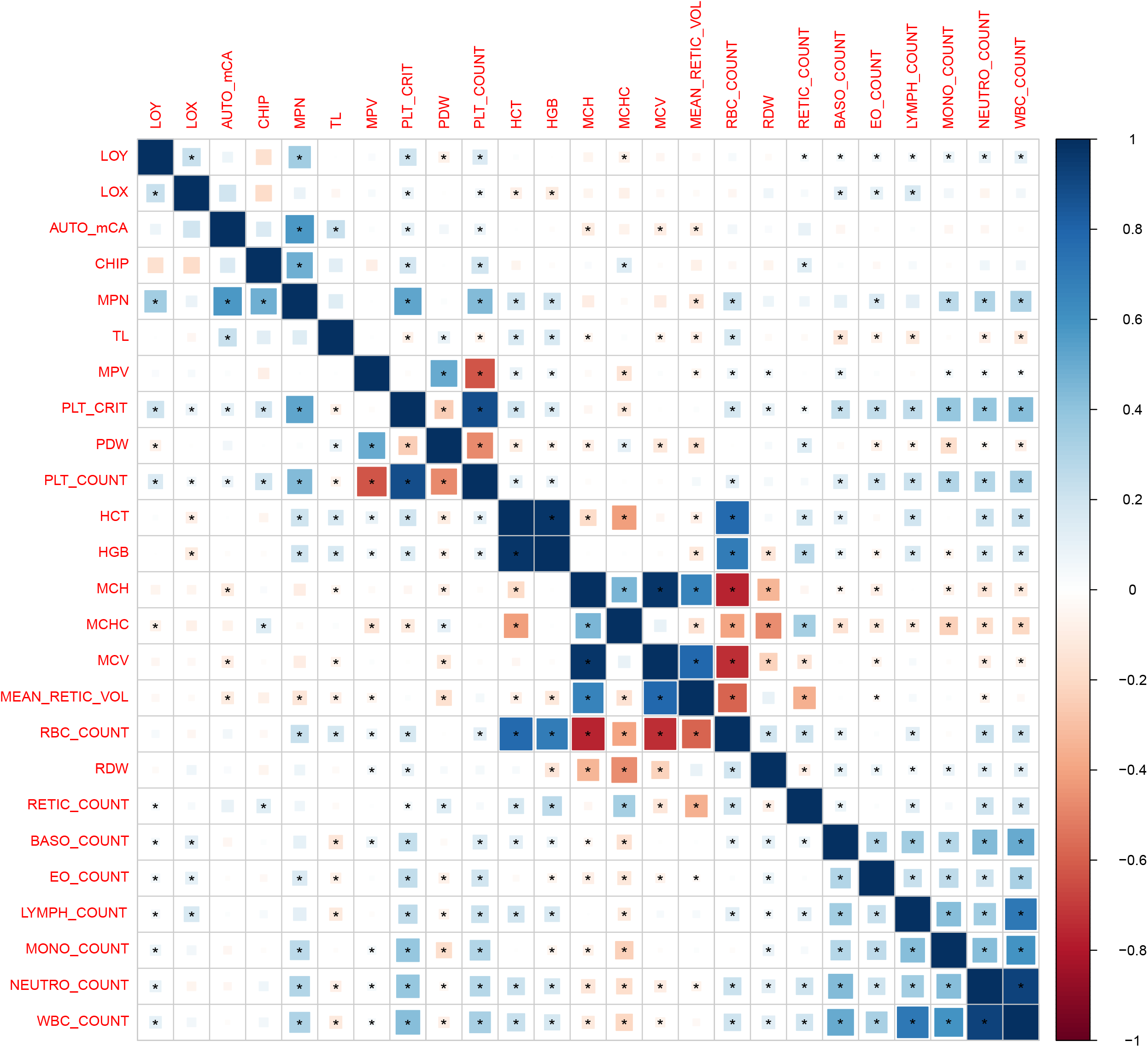
Pairwise genetic correlations between each type of clonal hematopoiesis, telomere length, and 19 blood cell traits derived using the high-definition likelihood (HDL) method. Square areas represent the absolute value of genetic correlations. Blue, positive genetic correlation; red, negative genetic correlation. Genetic correlations that are significantly different from zero (p-value< 0.05) are marked with an asterisk. All pairwise genetic correlations and p-values are given in **Supplemental Table 1**.

In an analysis of 482,378 subjects from the UK Biobank, we investigated adjusted phenotypic associations between types of CH (**Online Methods, Supplemental Table 3**). Consistent with previous studies of CH,^12–14^ each type of CH investigated demonstrated a strong positive association with age (**Supplemental Figure 2**). We observed an inverse phenotypic association between LOY and MPN (T-statistic= -4.76, P= 1.96×10^−6^) (**Figure 3, Supplemental Table 4**), which is opposite in direction from the genetic correlation. We were unable to evaluate the phenotypic association between LOY and LOX, as these are sex specific traits.

**Figure 3.**
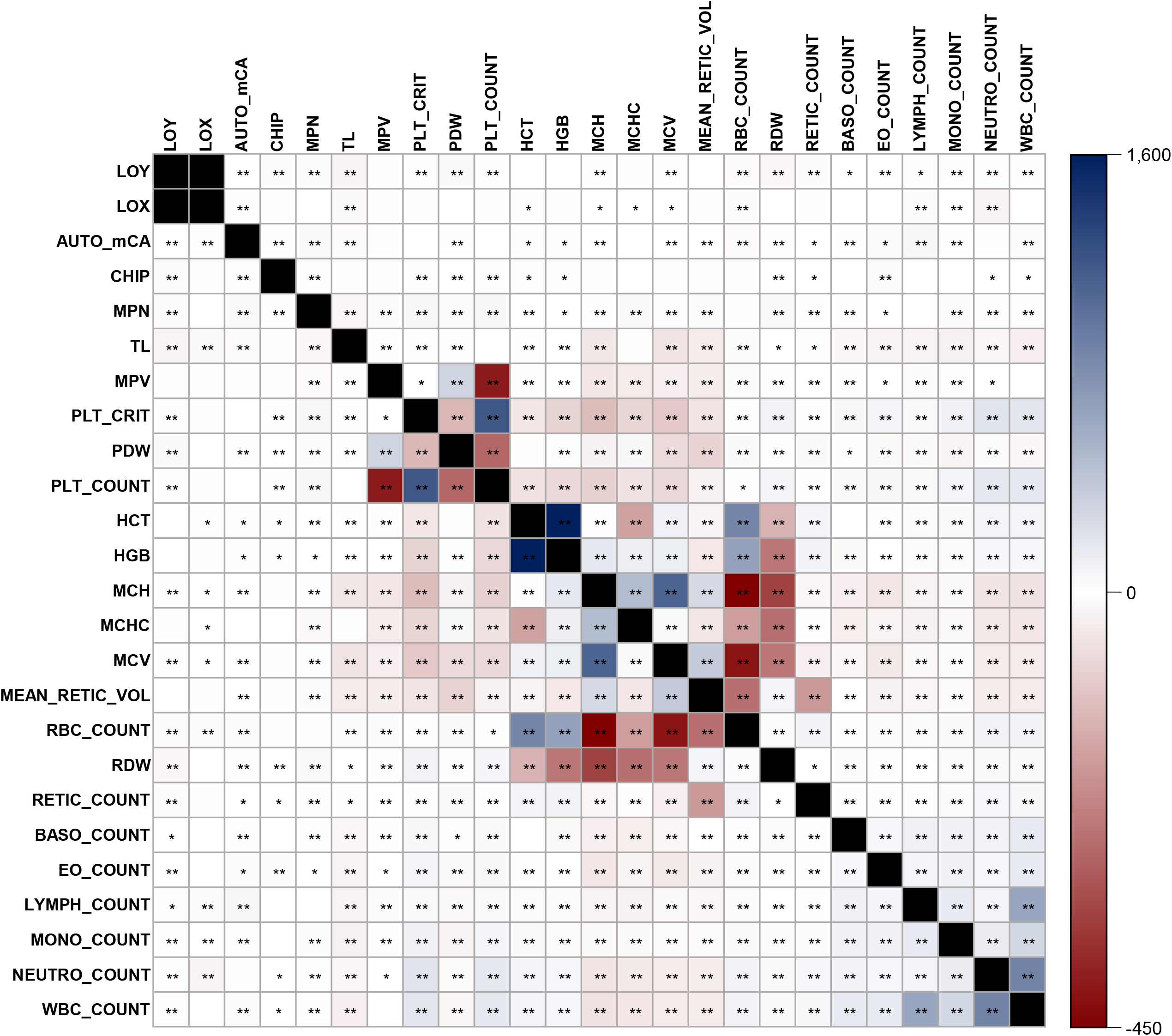
Pairwise phenotypic associations between each type of clonal hematopoiesis, telomere length, and 19 blood cell traits. Blue, positive T-statistic; red, negative T-statistic. T-statistics were derived using linear regression adjusted for age, age-squared, 25-level smoking status, and sex (in non LOY or LOX comparisons). Black cells were not tested. T-statistics that are significantly different from zero at a nominal p-value (p< 0.05) are marked with an asterisk and Bonferroni corrected p-value (p< 1.67×10^−4^) are marked with two asterisks. All pairwise T-statistics and p-values are given in **Supplemental Table 4**.

We report positive phenotypic associations of autosomal mCAs with LOY (T-statistic= 4.31, P= 1.61×10^−5^), LOX (T-statistic= 12.44, P= 1.54×10^−35^), CHIP (T-statistic= 8.89, P= 6.41×10^−19^), and MPN (T-statistic= 41.00, P< 5×10^−324^) (**Figure 3, Supplemental Table 4**). Sensitivity analyses removed individuals with mCAs spanning the *JAK2* region (N=550), a region frequently impacted by mCAs in MPNs,^15–18^ and still observed a positive phenotypic association between autosomal mCAs and MPN, though the association was attenuated (T-statistic= 15.53, P= 2.39×10^−54^). CHIP was also positively associated with MPN (T-statistic= 8.82, P= 1.18×10^−18^) and inversely associated with LOY (T-statistic= -4.11, P= 4.04×10^−5^) (**Figure 3, Supplemental Table 4**). The inverse association and exclusivity between CHIP and LOY was consistently observed when stratified by frequently observed CHIP gene mutations, *e,g*., *DNMT3A* CHIP with LOY (N_CHIP_= 1,818, T-statistic= -3.71, P= 2.08×10^−4^) and *TET2* CHIP with LOY (N_CHIP_= 786, T-statistic= -3.99, P= 6.50×10^−5^) (**Supplemental Table 5**). In further evaluation, we performed exome-wide burden analyses to identify rare (MAF<0.1%) protein coding variants associated with LOY (**Online Methods**). These analyses identified three established CHIP genes at exome-wide significance (**Supplemental Table 6**), demonstrating that individuals carrying heterozygous loss-of-function variants in *TET2* (n=193, beta = -0.21, SE = 0.03, P = 7.7×10^−15^), *ASXL1* (n=213, beta = -0.18, SE = 0.03, P = 1.3×10^−12^), and *DNMT3A* (n=89, beta = -0.17, SE = 0.04, P = 2.2×10^−5^) were less likely to exhibit LOY (**Supplemental Figures 3** and **4, Supplemental Table 7**). These findings reinforce the idea that acquiring LOY in the presence of CHIP mutations is likely selected against in clonally-expanded hematopoietic stem cells.

We next examined the cellular fraction of individuals with autosomal mCA events and the variant allele frequency (VAF) of individuals with CHIP mutations and observed that individuals with higher cellular fractions of autosomal mCA events (*i*.*e*., greater proportion of cells carrying the somatic event) were more likely to have LOX (T-statistic= 3.08, P= 2.09×10^−3^), CHIP (T-statistic= 2.35, P= 1.88×10^−2^), and MPN (T-statistic= 17.83, P= 3.17×10^−70^) (**Supplemental Figure 5, Supplemental Table 8**). Higher autosomal mCA cellular fraction was inversely associated with LOY (T-statistic= -7.76, P= 9.77×10^−15^) (**Supplemental Figure 5, Supplemental Table 8**). Individuals with higher VAF of CHIP mutations (*i*.*e*., higher clonal fractions) were more likely to have detectable autosomal mCAs (T-statistic= 5.82, P= 6.21×10^−9^) and MPNs (T-statistic= 7.19, P= 7.36×10^−13^), and less likely to have LOY (T-statistic= -3.80, P= 1.48×10^−4^) (**Supplemental Figure 6, Supplemental Table 9**).

In an analysis of co-existence of types of CH, CHIP and autosomal mCAs significantly co-occurred in the same individual (hypergeometric P= 5.32×10^−28^; **Supplemental Figure 7a**) with 439 individuals (6.0% of individuals with CHIP, 6.3% of individuals with autosomal mCAs) carrying both (**Supplemental Table 3**). Individuals with autosomal mCAs displayed a distinct pattern of CHIP gene mutations compared to individuals without autosomal mCAs (**Supplemental Figure 7b, Supplemental Table 10**). 13 CHIP gene mutations were significantly enriched in individuals with autosomal mCAs (*DNMT3A, TET2, ASXL1, TP53, SF3B1, STAT3, SRSF2, MPL, KRAS, JAK2, IDH1, PRPF40B*, and *PIGA*; **Supplemental Figure 7b**), a similar pattern of co-occurrence as previously observed.^10,11^ Additionally, individuals with CHIP mutations were more likely to acquire autosomal mCAs across 16 chromosomes (**Supplemental Figure 7c, Supplemental Table 11**), with enrichment for several chromosome-specific copy number states (*e*.*g*., CNLOH in chromosomes 1, 4, and 9; **Supplemental Figure 7c, Supplemental Table 11**).

An evaluation of the 439 individuals with both a CHIP mutation and autosomal mCA revealed that 53 (12.1%) had events spanning the same genomic region (binomial P= 1.70×10^−10^). 9 CHIP genes overlapped with autosomal mCAs, with *TET2* mutations accounting for 34 (54.0%) of the observed overlapping mutations (**Supplemental Figure 8**). CNLOH was the most frequently observed autosomal mCA event (N= 46 (73.0%)) among all overlapping mutations (**Supplemental Figure 8**). We examined the clonal fractions of both somatic mutations to provide a window into the clonal evolution of CHIP mutations and autosomal mCAs and found higher estimated CHIP VAF than estimated mCA cellular fraction in a majority of co-localizing mutations, suggesting the acquisition of the CHIP mutation preceded the acquisition of autosomal mCAs (binomial P=1.75×10^−4^; **Supplemental Figure 9**); this finding is consistent with a multi-hit hypothesis in driving clonal evolution. This is particularly evident in loss-of-heterozygosity of chromosome 9 alterations after acquisition of a *JAK2 V617F* mutation, as has been seen in individuals with MPNs.^19,20^ Subsequent autosomal mCA-induced loss of heterozygosity or amplification of CHIP driver mutations could confer strong selective advantages promoting rapid cellular expansion. To test this hypothesis, we investigated hematological malignancy risk in individuals with and without CHIP and autosomal mCAs (**Supplemental Figure 10**). Individuals with both CHIP and non-overlapping autosomal mCAs (N= 386) demonstrated a strong positive association with incident hematological malignancy risk (hazard ratio (HR) = 7.25, 95% confidence interval (CI) = 5.27-9.96, P= 2.94×10^−34^) compared to individuals without CHIP or autosomal mCAs. Individuals carrying overlapping CHIP and autosomal mCAs (N= 53) displayed an even stronger association (HR= 17.31, 95% CI= 9.80-30.58, P= 8.94×10^−23^), significantly elevated compared to individuals with non-overlapping CHIP and autosomal mCAs (P_heterogeneity_= 8.83×10^−3^, **Supplemental Figure 10** and **Supplemental Table 12**). The co-occurrence and overlap of CHIP and autosomal mCAs motivates future studies that jointly assess both CH traits to better understand CH interactions that could confer increased propensity for clonal expansion and elevated disease and mortality risk, particularly at specific loci or with specific mutations.^9^

Pathway-based analyses using GWAS summary statistics (**Online Methods**) utilized 6,290 curated gene sets and canonical pathways from Gene Set Enrichment Analysis (GSEA) and revealed significant associations between several biological pathways and types of CH (**Supplemental Figure 11**) with all types of CH associated with gene sets related to apoptosis, IL-2 signaling, DNA methylation, promyelocytic leukemia gene product (PML) targets, and cancer-related gene sets (**Supplemental Tables 13**-**17**). LOY, LOX, and MPN were significantly associated with hematopoietic progenitor cells, hematopoietic cell lineage and differentiation gene sets, and DNA damage response (**Supplemental Tables 13, 14**, and **17**). LOY, autosomal mCAs, CHIP, and MPN were associated with telomere extension by telomerase, with LOY and MPN also associated with telomere stress induced senescence (**Supplemental Tables 13, 15-17**). Additionally, the 12 genetic pathways significantly associated with autosomal mCAs were also associated with CHIP, providing further evidence that these types of CH are interrelated. Overall, pathway analyses suggest core shared pathogenic mechanisms related to cellular differentiation, DNA damage repair, and cell cycle regulation that are critical for the development and clonal expansion of most types of CH.

### Correlation of types of CH with myeloid and lymphoid cell traits

We examined genetic and phenotypic correlations between types of CH and 19 blood cell traits to assess lineage-specific effects by type of CH (**Figures 2**-**3**). All types of CH displayed positive genetic correlations for both plateletcrit (P< 0.02) and platelet count (P< 0.05) (**Figure 2, Supplemental Table 1)**. LOY, MPN, and CHIP were the only types of CH to also display significant phenotypic associations with plateletcrit (P< 2×10^−13^) and platelet count (P< 1.5×10^−13^) (**Figure 3, Supplemental Table 4**). LOY and MPN demonstrated additional genetic correlations enriched for myeloid lineage traits, namely, positive correlations with total white blood cell, eosinophil, monocyte, and neutrophil counts (P< 0.026; **Figure 2, Supplemental Table 1**). LOY was additionally positively correlated with lymphocyte count (ρ= 0.05, P= 8.74×10^−3^; **Figure 2, Supplemental Table 1**). MPN was positively correlated with other myeloid lineage traits including hematocrit, hemoglobin, and red blood cell count (P< 6.5×10^−4^; **Figure 2, Supplemental Table 1**), as previously reported.^21^ In support of the genetic correlations, we observed strong phenotypic associations of LOY and MPN with myeloid traits that closely mirror the magnitude and significance of the genetic correlation results (**Figure 3, Supplemental Table 4**), and previously reported phenotypic associations.^22–24^ Both LOY and MPN were positively associated with monocyte, neutrophil, and white blood cell counts (P< 1.5×10^−41^; **Figure 3, Supplemental Table 4**). LOY was also inversely associated with lymphocyte count (T-statistic= -2.75, P= 5.91×10^−3^) (**Figure 3, Supplemental Table 4**). These findings suggest shared mechanisms regulating hematopoiesis likely also govern susceptibility to LOY and MPN.

LOX was the only type of CH to display both a positive genetic correlation (ρ= 0.17, P= 8.40×10^−5^; **Figure 2, Supplemental Table 1**) and a positive phenotypic association with lymphocyte count (T-statistic= 23.96, P= 9.11×10^−127^) (**Figure 3, Supplemental Table 4**). LOX had a positive genetic correlation with myeloid traits such as basophil count and eosinophil count, whereas it displayed an inverse genetic correlation with hematocrit and hemoglobin (P< 0.02, **Figure 2, Supplemental Table 1**). LOX also had positive phenotypic associations with MCH, MCHC, MCV, and monocyte count (P< 0.015, **Figure 3, Supplemental Table 4**), and inverse associations with hematocrit, red blood cell count, and neutrophil count (P< 0.03, **Figure 3, Supplemental Table 4**).

Besides the aforementioned genetic correlations with plateletcrit and platelet count, we observed additional genetic correlations between autosomal mCAs and CHIP with blood cell traits (**Figure 2**). Inverse genetic correlations were observed between autosomal mCAs with MCH, MCV, and mean reticulocyte volume (P< 3.0×10^−2^, **Figure 2, Supplemental Table 1**). CHIP had a positive genetic correlation with MCHC and reticulocyte count (P< 4.0×10^−2^, **Figure 2, Supplemental Table 1**). In the case of combined autosomal mCAs, there was evidence for positive phenotypic associations with both lymphocyte count (T-statistic= 60.33, P< 5×10^−324^) and total white blood cell count (T-statistic= 34.48, P= 3.89×10^−260^) (**Figure 3, Supplemental Table 4**). Combined CHIP was positively associated with platelet distribution width (T-statistic= 5.02, P= 5.26×10^−7^), red blood cell distribution width (T-statistic= 4.09, P=4.37×10^−5^), and neutrophil count (T-statistic= 3.59, P= 3.32×10^−4^) (**Figure 3, Supplemental Table 4**), all of which are myeloid lineage traits. The CHIP phenotypic association findings support recent evidence suggesting CHIP primarily results in myeloid-related disruptions, although select distinct CHIP events could increase risk for disruptions in the lymphoid lineage.^9^ Together our results support lineage-specific effects that differ by type of CH, suggesting shared etiology, specifically shared genetic etiology for LOY and myeloid traits, as well as ample phenotypic associations that detail early downstream phenotypic disruptions in hematologic phenotypes that alter disease risk.

### A dynamic association of telomere length with CH

Telomere length in leukocytes provides a metric of hematopoietic stem cell activity and can provide insights into how genetic variation in hematopoietic stem cells interact with risk for acquiring CH.^21,25^ The genetic relationship between each type of CH with leukocyte telomere length (TL) was evaluated to determine whether genetic variation in telomere maintenance genes could also contribute to predisposition to CH. A positive genetic correlation for autosomal mCAs with TL was observed (ρ= 0.23, P= 4.95×10^−3^) (**Figure 2, Supplemental Table 1**). To further test for a genetic association, we conducted one-direction Mendelian randomization (MR) between TL and each CH type using 130 previously published TL-associated variants (**Supplemental Figure 12**).^26^ Based on MR-IVW models, we observed positive associations between the TL IV and autosomal mCAs (Z_filtered_= 5.65, P= 1.21×10^−7^), CHIP (Z_filtered_= 5.72, P= 9.65×10^−8^), and MPNs (Z_filtered_= 5.61, P= 1.88×10^−7^), and observed a negative association between the TL IV and LOY (Z_filtered_= -6.40, P= 8.11×10^−9^) and did not identify evidence for a genetic relationship between telomere length and LOX (**Supplemental Figure 13, Supplemental Table 18**). These observations provide additional support of associations between inherited telomere length and select CH traits.^12,21,27–30^ The intercept from MR-Egger regression was significant (p< 0.05) for both autosomal mCAs and MPN (**Supplemental Table 18**), so we performed additional MR weighted median (MR-WM) analyses which displayed the same positive association between the TL IV and autosomal mCAs (Z_filtered_= 4.16, P= 6.19×10^−5^), and MPNs (Z_filtered_= 4.34, P= 3.53×10^−5^) (**Supplemental Table 18**). These MR associations are supported by our pathway analyses, which demonstrate telomere pathways are significantly associated with LOY, autosomal mCAs, CHIP, and MPN (**Supplemental Tables 13, 15-17**). Based on these data it is plausible that inherited variation in telomere length maintenance contributes to clonal expansion of mutated hematopoietic stem cells, or alternatively confers greater risk for mutation acquisition and clonal evolution in hematopoietic stem cells.

Once clonal expansion ensues, measured telomere length is a metric of hematopoietic stem cell growth and clonal expansion. Using available measured telomere length data from UK Biobank, we observed inverse phenotypic associations between CH and measured telomere length (**Figure 3**). CHIP, which presents with the smallest fraction of mutated clones, had an insignificant phenotypic association with measured TL (T-statistic= -1.03, P= 0.30) (**Figure 3, Supplemental Table 4**). To further examine this relationship, we conducted analyses between CHIP VAF and measured TL and observed individuals with higher VAF, *i*.*e*., higher CHIP cellular fraction, had a more inverse association with measured TL (T-statistic= -6.50, P= 8.34×10^−11^, **Supplemental Figure 14a** and **Supplemental Table 19**). Additionally, individuals with higher autosomal mCA cellular fraction also demonstrated a stronger inverse association with measured TL (T-statistic= -9.02, P= 2.16×10^−19^, **Supplemental Figure 14b** and **Supplemental Table 20**). The number of mutations present in an individual was also inversely associated with TL for increasing autosomal mCA count (T-statistic= -10.01, P= 1.35×10^−23^). Individuals with both CHIP and autosomal mCAs also demonstrated an inverse association with TL (T-statistic= -2.75, P= 5.93×10^−3^) with individuals carrying overlapping CHIP and autosomal mCAs displaying a stronger inverse association with TL (T-statistic= -3.48, P= 5.01×10^−4^) compared to individuals without CHIP or autosomal mCAs (**Supplemental Figure 15** and **Supplemental Table 21**). These inverse TL associations indicate increased clonal expansion leads to reduced measured telomere length and suggest reductions in telomere length from the expansion of mutated clones could lead to further genomic instability and the acquisition of additional CH mutations.

### Leveraging shared correlations to nominate additional MPN susceptibility loci

Finally, we leveraged the shared genetic architecture between these CH traits (**Figure 4**) to identify novel loci associated with MPN - a disease where it has been challenging for GWAS to perform well powered case-control analyses, despite the finding of considerable heritable influences on this disorder.^21^ We first performed multi-trait analysis of GWAS (MTAG), which boosts the power to identify potential MPN-associated signals by leveraging the shared genetic architecture with LOY and TL (**Online Methods**). This approach identified 25 MPN loci at genome-wide significance (P<5×10^−8^), 15 of which have not been previously implicated in MPN (**Supplemental Table 22**). We next evaluated a complementary approach of performing colocalization analyses (**Online Methods**) using genome-wide significant loci associated with LOY, TL, and MPN. We found that 12 LOY loci, mapping to 11 genes (*TET2, NREP, GFI1B, TERT, DLK1, PARP1, TP53, RBPMS, MAD1L1, MECOM*, and *ATM*) co-localized with MPN (**Supplemental Table 23**), highlighting 6 loci that have not previously reached genome-wide significance for MPN (P= 1.17×10^−4^ to 5.14×10^−8^). In addition, 5 leading SNPs for TL co-localized with MPN and mapped to 4 genes (*TERT, NFE2, PARP1*, and *ATM*), 2 of which have not reached genome-wide significance (P<5×10^−8^) in prior MPN analyses (*NFE2* and *PARP1*) (**Supplemental Table 24**). Of note, leading SNPs at *TERT, PARP1*, and *ATM* colocalized across all 3 traits (**Supplemental Table 23** and **Supplemental Table 24**), and 5 co-localized loci also reached genome-wide significance in the MTAG analysis (*PARP1, MAD1L1, DLK1, RBPMS*, and *TP53*) (**Figure 5**). While validation is required for the newly identified putative MPN risk loci, these results illuminate opportunities to use insights from correlated diseases or phenotypes to gain new genetic and biological insights on blood cancer risk.

**Figure 4.**
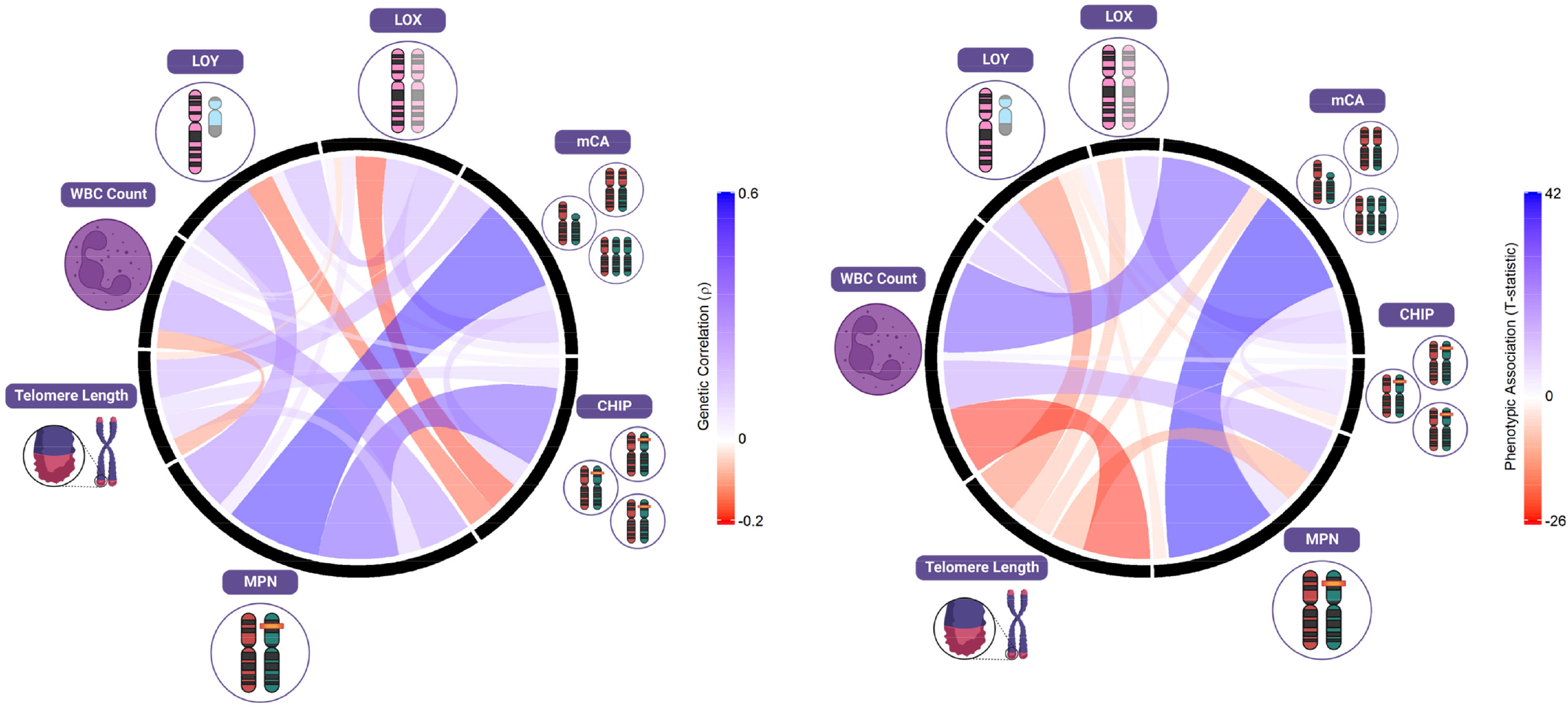
Shared etiologies and associations between types of CH and hematopoietic phenotypes: telomere length, and white blood cell count (WBC). Pairwise HDL genetic correlations are given in the left plot, pairwise phenotypic associations derived using linear regression adjusted for age, age-squared, 25-level smoking status, and sex (in non LOY or LOX comparisons) are given in the right plot.

**Figure 5.**
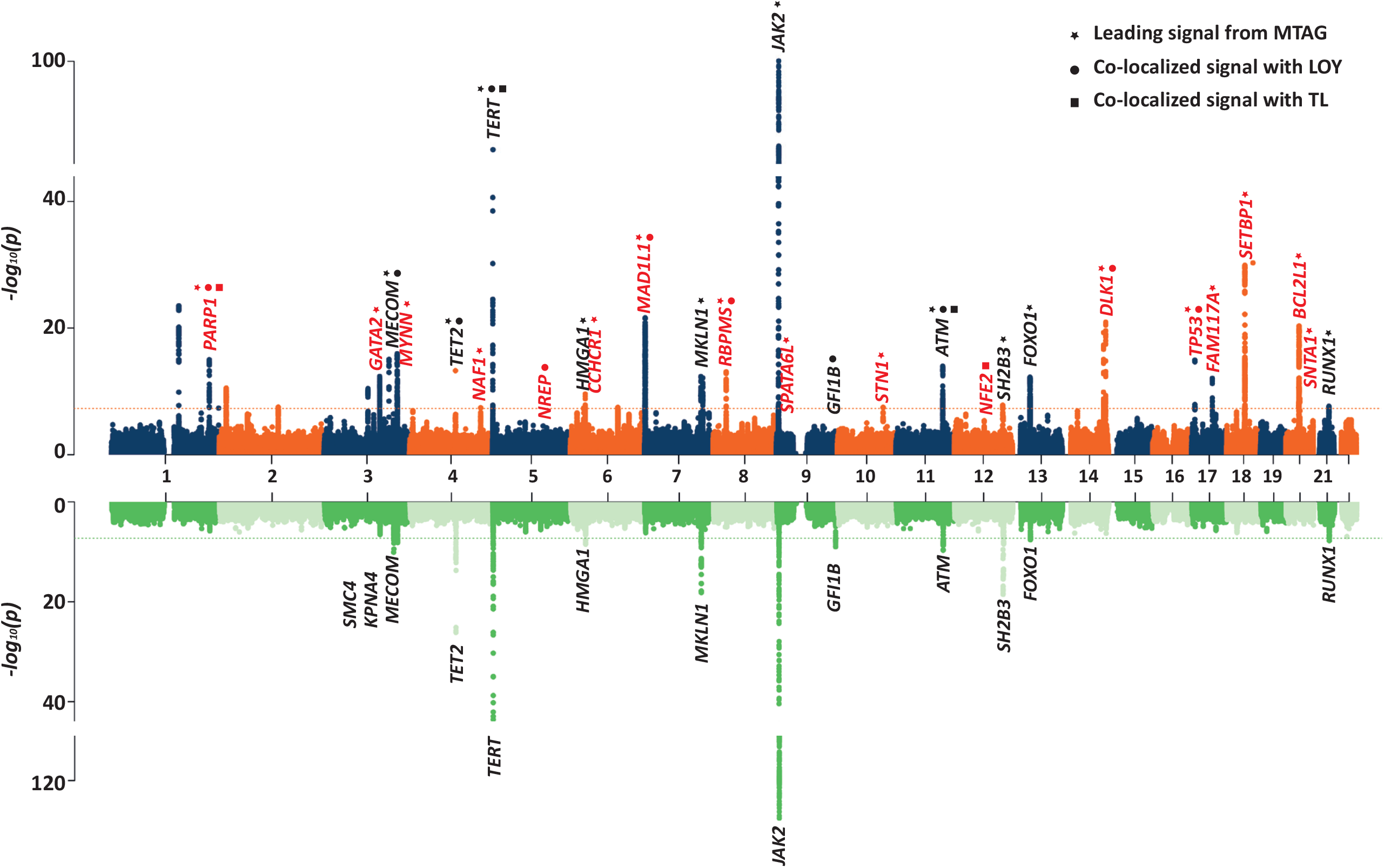
Stacked Manhattan plots from the MTAG and colocalization analyses among telomere length, LOY, and MPN (Top plot) and the original MPN GWAS performed by Bao et al. (Bottom plot). Nominated MPN susceptibility loci are labeled by analysis, MTAG((⍰), LOY colocalization(•), TL colocalization(⍰), and colored red for previously unidentified and black for previous identified.

## Discussion

Understanding the underlying molecular mechanisms of different types of CH is critical for disentangling age-related clonal evolution and the possible impact of CH on subsequent disease risk, particularly the risk of acquiring blood cancers. Herein, our analysis of CH using large-scale genetic data highlights both similarities in the underlying mechanisms and key differences, particularly with respect to events related to distinct aspects of hematopoiesis. Common to the types of CH are core pathways, namely, cellular differentiation, DNA repair, and cell cycle regulation, that contribute to generation and clonal expansion. Together, these findings detail specific characteristics of CH that should be investigated to improve the utility of detectable CH for disease risk and possible intervention or prevention.

We provide evidence for genome-wide genetic correlations between LOY and LOX, LOY and MPN, autosomal mCAs and MPN, as well as between CHIP and MPN, suggesting shared biologic mechanisms promoting or predisposing to the development and clonal expansion. Genetic correlations with blood cell traits further demonstrate lineage-specific effects that differ by type of CH. As many of these CH and blood cell traits are interrelated, we report associations that do not adjust for stringent multiple testing corrections and caution against the overinterpretation of marginally significant associations. Phenotypic associations between types of CH further support similar genetic etiology, but could also indicate shared environmental factors that drive CH growth and expansion (*e*.*g*., medication^31,32^ or tobacco usage^12,33–37^). A notable discordance in directionality between the genetic correlation and the phenotypic relationship between LOY and MPN supports a shared genetic etiology, but also suggests a mutual exclusivity of some CH types such that once one type of CH develops, the occurrence of others may be suppressed (*e*.*g*., *DNMT3A* and *TET2* CHIP^38^). Individuals with CHIP and higher cellular fraction autosomal mCA events also demonstrated an inverse phenotypic association with LOY indicating a similar mutually exclusive relationship. These findings further support common genetic factors, and raise the importance of pursuing shared environmental contributors beyond smoking and air-pollution.^12,33–37,39^ Our findings reveal that there is mutual exclusivity of CH, most likely due to hematopoietic stem cells that cannot tolerate multiple independent somatic drivers of CH. However, we also observe strong evidence for the co-occurrence of CHIP and autosomal mCAs in the same individual, and in many instances, overlapping within known CH driver mutations (*e*.*g*., *TET2, DNMT3A, JAK2*). Cross-sectional observations of cellular fraction indicate these CHIP mutations often precede autosomal mCAs, which can lead to preferential clonal expansion of mCAs containing CHIP mutations, as has been mechanistically examined in specific cases.^40^

Rapid clonal expansion afforded by each type of CH leads to marked reductions in measured telomere length. These reductions in telomere length can lead to increased genomic instability in individuals with CH and could increase the likelihood of acquiring additional types of CH. Individuals who acquired overlapping CHIP and autosomal mCAs were found to have greater reductions in telomere length, which is a marker of past clonal expansion, while also having a significantly higher risk for acquiring blood cancers. Detection of these highly clonal, co-occurring CH events, especially at *TET2, DNMT3A* and *JAK2*, could be helpful in identifying individuals at increased risk of developing hematological malignancies. Likewise, these genetic relationships can be leveraged to identify disease susceptibility loci for related traits. Future studies should focus on investigating the co-occurrence and overlap of CHIP and autosomal mCAs to further evaluate associations with environmental factors, elevated disease, and mortality.

## Online Methods

### Hematopoietic Phenotypes

We used genome-wide association study (GWAS) summary statistics to investigate germline similarities and differences of 25 hematopoietic related phenotypes. These included loss of chromosome Y (LOY) in men,^13^ loss of chromosome X (LOX) in women, autosomal mCAs including gains, losses and copy neutral loss of heterozygosity,^29^ CHIP, MPNs,^21^ leukocyte telomere length (TL),^26^ and 19 blood cell traits^21^ (**Supplemental Table 25**). For LOX, we used previously generated data on copy number variation,^24,29^ and performed GWAS on 243,106 women in the UK Biobank using a linear mixed model implemented in BOLT-LMM,^41^ to account for cryptic population structure and relatedness, a similar methodology was used to conduct the LOY GWAS.^13^ For CHIP, we called somatic mutations using Mutect2 from available UK Biobank 200K whole exome sequencing data.^42^ A QCed set of N=198,178 individuals were analyzed using a panel of one hundred normals created from UK Biobank participants with age <40, and included as part of the QC process. Variants were considered passing QC if the following criteria were met: meeting FilterMutectCalls quality standards including learned read orientation, variant allele fraction (VAF) >=2%, depth of calling >10, and a Phred scaled GERMQ score of >20 (1% error rate). CHIP was defined in these individuals using a curated list of CHIP mutated variants as previously described in the UKBB WES cohort (**Supplemental Table 26**).^43^ Individuals with a diagnosis of myeloid malignancy (AML, MDS, MPN) before blood draw were excluded from the CHIP phenotype while those that went on to develop myeloid malignancy post-blood draw by at least 5 years were retained. In total, 7,280 (3.7%) individuals were found to have at least one CHIP curated variant (**Supplemental Table 26, Supplemental Table 27**). We performed a CHIP GWAS in the UK Biobank array data,^44^ restricted to European ancestry individuals (**Supplemental Table 25**) and those passing the following QC measures: individual had not withdrawn consent, included in kinship inference, no excess (>10) of putative third-degree relatives inferred from kinship, not an outlier in heterozygosity and missing rates, not found to have putative sex chromosome aneuploidy, no genotype missing rate of >0.1. Variants were included if they had a genotype missing rate <0.1 across QC’ed individuals, Hardy-Weinberg equilibrium p-value of >1×10^−15^, and a minor allele frequency of >0.1%. Using Regenie,^44^ we performed ridge regression in step 1 using a set of ∼300,000 pruned SNPs and default cross-validation settings. We included age, age-squared, sex, smoking status (categorical level variable), and principal components of ancestry 1 through 10 as covariates. To further increase the power of downstream analyses, we conducted a meta-analysis using METAL^45^ of CHIP GWAS summary statistics between our generated UK Biobank summary statistics and those from a previous CHIP GWAS in TOPMed.^12^ Before meta-analysis, both GWAS summary statistics were lifted-over from their respective genome builds to reference genome hg19. Alleles were flipped according to the hg19 build reference allele, and if neither allele was present the variant was removed. Strand ambiguous and non-biallelic SNPs were removed. Minor allele frequency was filtered to >=0.1%. RSIDs were assigned to variants using dbSNP version 144.

Data on 482,378 subjects from the UK Biobank were used for phenotypic association analyses, after removing individuals with sex discordance or whose DNA failed genotyping QC. We used previously generated data for each of the hematopoietic related phenotypes,^13,21,26,29^ with the exception of CHIP which was called in the UK Biobank as detailed above (**Supplemental Table 3**).

### Genetic Correlation

We used both the high-definition likelihood method^46^ (HDL) and linkage disequilibrium score regression^47^ (LDSC) to compute pairwise genetic correlation between hematopoietic phenotypes. For HDL, we utilized an LD score reference panel available within HDL which contains 1,029,876 QCed UK Biobank imputed HapMap3 SNPs,^46^ and also calculated the observed heritability for each hematopoietic phenotype (**Supplemental Figure 16** and **Supplemental Table 28**). For LDSC, we utilized an LD score reference panel generated on 6,285 European ancestry individuals combined from the 1000 Genomes Phase 3 and UK10K cohorts, with a total of 17,478,437 available variants, and GWAS summary statistics were filtered to include overlapping variants with >1% MAF and >90% imputation quality score. As all of the included GWAS summary statistics utilized UK Biobank data, we constrained the intercept within both HDL and LDSC by accounting for both the known sample overlap and phenotypic correlation between traits (**Supplemental Table 29**). Pairwise genetic correlations with CHIP were conducted using the generated UK Biobank summary statistics, due to incomplete overlap with the HDL reference panel within the TOPMed CHIP summary statistics (**Supplemental Table 25** and **Supplemental Table 29**). Correlation matrixes and circular charts for results visualization were generated using the “corrplot” and “circlize” R packages.^48,49^

### Phenotypic Associations

Pairwise phenotypic associations between all hematopoietic phenotypes were generated using linear regression adjusting for age, age-squared, sex (in non sex-specific traits), and a 25-level smoking variable to reduce the potential for confounding variables driving associations.^35^ To ensure compatibility between binary and continuous phenotypic association results, association T-statistics were generated and reported to measure strength and direction of phenotypic associations.

### Hematological Malignancy Association

Using available data within UK Biobank, we extracted relevant cancer information from both inpatient records and cancer registry data. Incident hematological cancers were defined as occurring after study enrollment using codes: C81: Hodgkin’s disease, C82: Follicular [nodular] non-Hodgkin’s lymphoma, C83: Diffuse non-Hodgkin’s lymphoma, C84: Peripheral and cutaneous T-cell lymphomas, C85: Other and unspecified types of non-Hodgkin’s lymphoma, C86: Other specified types of T/NK-cell lymphoma, C88: Malignant immunoproliferative diseases, C90: Multiple myeloma and malignant plasma cell neoplasms, C91: Lymphoid leukemia, C92: Myeloid leukemia, C93: Monocytic leukemia, C94: Other leukaemias of specified cell type, C95: Leukemia of unspecified cell type, C96: Other and unspecified malignant neoplasms of lymphoid, haematopoietic and related tissue, D47: Other neoplasms of uncertain or unknown behavior of lymphoid, haematopoietic and related tissue. We performed Cox proportional hazards regression to assess the risk of hematological malignancies across CHIP and autosomal mCA group adjusting for age, age-squared, sex, and a 25-level smoking variable.^35^ Hazards ratios and 95% confidence intervals were generated and reported to measure strength and direction of hematological malignancy risk.

### Mendelian Randomization Analyses

We performed one-direction Mendelian randomization (MR) between TL and LOY, LOX, autosomal mCAs, CHIP, and MPN. Briefly, MR analyses utilize genetic variants from GWAS as instrumental variables (IVs) to assess the directional association between an exposure and outcome, which can mimic the biological link between exposure and outcome.^50^ Each variant used in a MR analysis must satisfy three assumptions: 1) it is associated with the risk factor, 2) it is not associated with any confounder of the risk factor–outcome association, 3) it is conditionally independent of the outcome given the risk factor and confounders.^51,52^

For our analyses, we used summary statistics and 130 significant signals of the largest TL GWAS to date to form the TL IV.^26^ We then extracted the same set of signals from summary statistics for each CH outcome. If any signals were missing in the outcome summary statistics, we collected proxies for these signals using GCTA^53^ with European UK Biobank individuals as reference (within 1 MB of reported signals and R^2^ > 0.4). We chose the proxy of each missing signal with the largest R^2^ value as the replacement IV, which was contained in both GWAS summary statistics of the exposure and outcome. All TL signals were aligned to increasing allele and alleles for outcome were realigned accordingly.

The MR inverse-variance weighted (MR-IVW) model, which can provide high statistical power,^54^ was used as our primary analysis. As some signals may have a stronger association with the outcome than the exposure, which may induce reverse causality, we applied Steiger filtering to each IV in order to remove these variants using the “TwoSampleMR” R package.^55^ We then applied Radial filtering to exclude signals that were identified as outliers according to Ru□cker’s Q′ statistic.^56^

The sensitivity of MR models was checked by the degree of heterogeneity (I^2^ statistics and Cochran’s Q-derived P-value), horizontal pleiotropy (MR-Egger p_intercept_ <0.05), and funnel and dosage plots (**Supplemental Figure 12**). To account for potential horizontal pleiotropy and heterogeneity, three additional MR models were performed: MR-Egger,^57^ weighted median (MR-WM),^58^ and penalized weighted median (MR-PWM).^58^

### Pathway and Gene Set Analyses

We performed agnostic pathway-based analyses using the summary data-based adaptive rank truncated product (sARTP) method, which combines GWAS summary statistics across SNPs in a gene or a pathway,^59^ to identify gene sets and pathways associated with each type of CH. A total of 6,290 curated gene sets and canonical pathways from GSEA (https://www.gsea-msigdb.org/gsea/msigdb/) were used for the analyses. For each type of CH, the signals from up to five of the most associated SNPs in a gene were accumulated. We adjusted for the number of SNPs in a gene and the number of genes in a pathway through a resampling procedure that controls for false positives.^59^ The P values of gene- and pathway-level associations were estimated from the resampled null distribution generated from 100 million permutations. Linkage disequilibrium between SNPs was computed from European individuals within 1000 Genomes Project data.^60^ To reduce the potential for population stratification to bias the results, we rescaled the marginal SNP results for each CH trait to set the genomic control inflation factor to 1. A Bonferroni corrected level of significance of 7.95×10^−6^ (0.05/6,290 GSEA pathways) was used to assess statistical significance.

### Rare variants gene-burden test for LOY in UK Biobank

To explore the relationship between rare variant burden and LOY, we performed association tests using whole exome sequencing (WES) data for 190,759 males provided by the UK Biobank. Prior to performing association tests, we performed quality control on provided sequencing data as previously described.^61^

We utilized the ENSEMBL Variant Effect Predictor (VEP) v104^62^ to annotate variants on the autosomal and X chromosomes. VEP was run with default settings, the “everything” flag, and the LOFTEE plugin.^63^ The predicted consequence of each variant was prioritized by a single MANE (version:0.97) or, when not available, a VEP canonical ENSEMBL transcript, and the most damaging consequence as defined by VEP defaults. Variants with high confidence (HC, as defined by LOFTEE) stop gained, splice donor/acceptor, and frameshift consequences were grouped as protein-truncating variants (PTVs). Following transcript annotation, we utilized CADD v1.6 to calculate the Combined Annotation Dependent Depletion (CADD) score for each variant.^64^

To perform gene burden tests, we implemented BOLT-LMM v2.3.6.^41^ As input, BOLT-LMM requires genotyping data for variants with allele count greater than 100, all variants from WES passing QC as defined above, and a set of dummy genotypes representing participant carrier status per-gene for PTVs, missense variants with CADD ≥ 25 (MISS_CADD25,) and damaging variants (HC_PTV+MISS_CADD25, DMG). Dummy genotypes were generated by collapsing all variants within each gene with a minor allele frequency (MAF) < 0.1%. For each gene, carriers with a qualifying variant were set to heterozygous (“0/1”) and non-carriers were set as homozygous reference (“0/0”). All models were controlled for age, age-squared, WES batch, and the first ten genetic ancestry principal components (PCs) as generated by Bycroft et al.^65^

Following association testing, we further excluded genes with less than 50 non-synonymous variant carriers, leaving 8,984 genes of PTVs, 14,685 genes of MISS_CADD25, and 16,066 genes of DMG for an exome-wide significance threshold of 1.26×10^−6^ (0.05/39,735) after Bonferroni correction (**Supplemental Table 6**).

### Associations between CHIP loss of function variant carriers and LOY

As associations between known CHIP genes and LOY identified as part of rare variant burden testing could be due to reverse-causality – somatic instability such as LOY could lead to, or occur in parallel with, variants arising within CHIP genes – we queried underlying variant call data to determine if individual variants within these genes were likely to have arisen somatically. We first extracted the number of reads supporting the alternate and reference alleles for all carriers of protein truncating variants (PTV) at MAF < 0.1% in four genes associated with LOY – three known CHIP genes, *ASXL1* (n = 213 carriers), *DNMT3A* (n = 89), and *TET2* (n = 193), and one control gene not previously associated with CHIP, *GIGYF1* (n = 81; **Supplemental Figure 3**). This information was then used to calculate a Variant Allele Fraction (VAF) for each genotype, where a VAF of 0.5 indicates perfect balance between sequencing reads supporting the reference and alternate allele (**Supplemental Figure 4**). For all variants, we also annotated whether it was found in a list of known, specific CHIP driver mutations or was likely to be a CHIP driver mutation based on a broader set of criteria presented in Bick et al.^12^ For each gene, we tested for an association between PTV carrier status and PAR-LOY except using 6 additional criteria that excluded individuals carrying:

1. Frameshift InDels with a binomial test p-value for allele balance < 0.001 (i.e. filtering InDels identically to SNVs, see above).
2. Any variant with VAF < 0.25 or > 0.75.
3. Any variant with VAF < 0.4 or > 0.6.
4. Any variant with VAF > 0.35.
5. A variant explicitly listed in Supplementary Table 3 from Bick et al.^12^
6. A variant explicitly listed in Supplementary Table 3 or matching the criteria in Supplementary Table 2 from Bick et al.^12^

All association tests were run separately for each gene using a logistic model corrected for identical covariates as the rare variant burden tests outlined above.

### MTAG and Colocalization analysis among TL, LOY, and MPN

GWAS summary statistics for LOY,^13^ TL,^26^ and MPN^21^ were utilized to conduct a meta-analysis by implementing the multi-trait analysis of GWAS (MTAG).^66,67^ Based on the summary statistics from GWAS of multiple correlated traits, MTAG can enhance the statistical power to identify genetic associations for each trait included in the analysis.^66,67^ We performed the MTAG analysis using Python. Prior to the analysis, we excluded the variants with MAF < 0.01 from the summary statistics of all three traits.^13,21,26^ A potential problem for MTAG is that SNPs can be null for one trait but non-null for another trait, which can cause MTAG’s effect size estimations of these SNPs for the first trait to shift away from 0. This causes the false positive rate (FDR) to increase. Therefore, we estimated the max FDR for each trait by invoking “—fdr” when running MTAG. We implemented a clumping algorithm to select signals from the MTAG generated MPN summary statistics. Preliminary leading signals were selected with a P< 5×10^−8^ and a MAF > 0.1% at a 1 Mb window. We then selected the secondary leading signals using approximate conditional analyses in GCTA^68^ with UK Biobank reference panel. If the genome-wide significant leading signals shared any correlation with each other due to the long-range linkage disequilibrium (r^2^> 0.05), these signals were excluded from further analysis. We mapped the leading signals to the genes with 1 Mb window based on the start and end sites of genes’ GRCh37 coordinates. For all leading signals, we extracted their summary statistics from the original MPN GWAS summary statistics. In total, 36 independent leading signals were identified. We then applied Bonferroni correction for the identified signals. We further excluded the signals with P > 0.05/36=1.39×10^−3^ in the original GWAS to avoid false positives mentioned above, as GWAS for both LOY and TL identified many more leading signals than MPN, which increased the FDR for MPN (Max FDR of MPN=0.11).

We conducted the Bayesian test for colocalization between pairs of TL and MPN, and LOY and MPN using their summary statistics^13,21,26^ and the leading GWAS signals by implementing R package coloc (Version: 5.1.0).^69^ The signals with posterior probability (h4.pp) ≥ 0.75 were defined as the co-localized causal variant for both traits. Manhattan plots for results visualization were generated using the “qqman” R package.^70^

## Acknowledgments

This work was conducted using the UK Biobank resource (applications 9905, 21552, and 31063). The UK Biobank was established by the Wellcome Trust, the Medical Research Council, the United Kingdom Department of Health, and the Scottish Government. The UK Biobank has also received funding from the Welsh Assembly Government, the British Heart Foundation, and Diabetes UK. The clonal hematopoiesis infographic and models were created with BioRender.com. We thank Dr. Jacqueline B. Vo for her graphical support. The opinions expressed by the authors are their own and this material should not be interpreted as representing the official viewpoint of the U.S. Department of Health and Human Services, the National Institutes of Health or the National Cancer Institute.

## Author contributions

J.R.B.P, V.G.S., and M.J.M. conceived the study. L.D.C. and V.G.S. carried out the CHIP calls. D.W.B., L.D.C., Y.Z., E.J.G., A.D., T.R., L.S., K.Y. performed computational and statistical analyses. J.R.B.P, V.G.S., and M.J.M. supervised the study. D.W.B., L.D.C., Y.Z., S.K.N., S.J.C., J.R.B.P, V.G.S., and M.J.M. drafted the manuscript with input from all authors. All authors critically read and approved the final version of the manuscript.

## Data Availability

All data used in analyses is available through application to the UK Biobank.

