## Supplemental Figures for "Shared and distinct genetic etiologies for different types of clonal hematopoiesis"

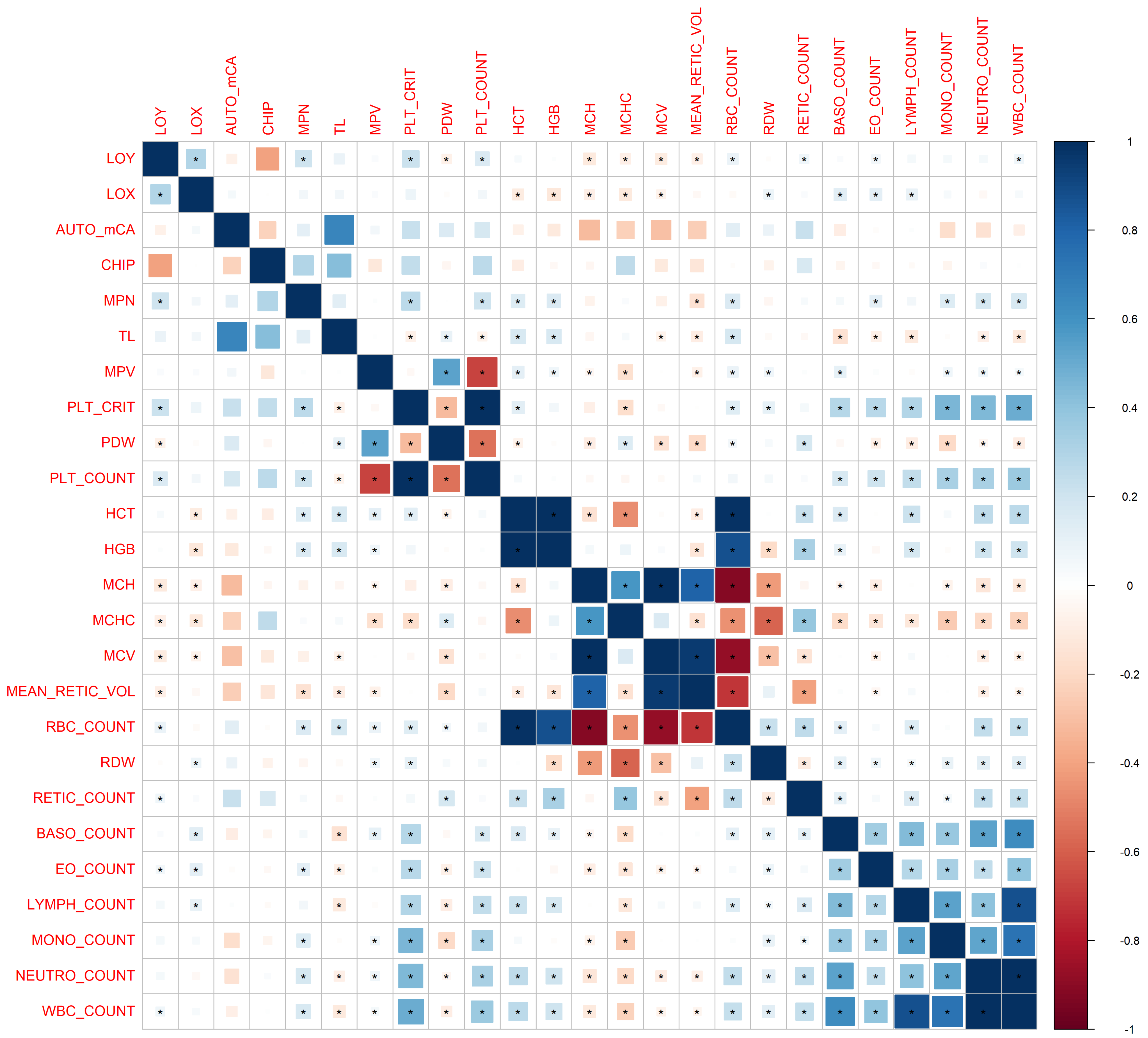


**Supplemental Figure 1**. Pairwise genetic correlations between each type of clonal hematopoiesis, telomere length, and 19 blood cell traits derived using linkage disequilibrium score regression (LDSC). Square areas represent the absolute value of genetic correlations. Blue, positive genetic correlation; red, negative genetic correlation. Genetic correlations that are significantly different from zero (p-value< 0.05) are marked with an asterisk. All pairwise genetic correlations and p-values are given in **Supplemental Table 2**.

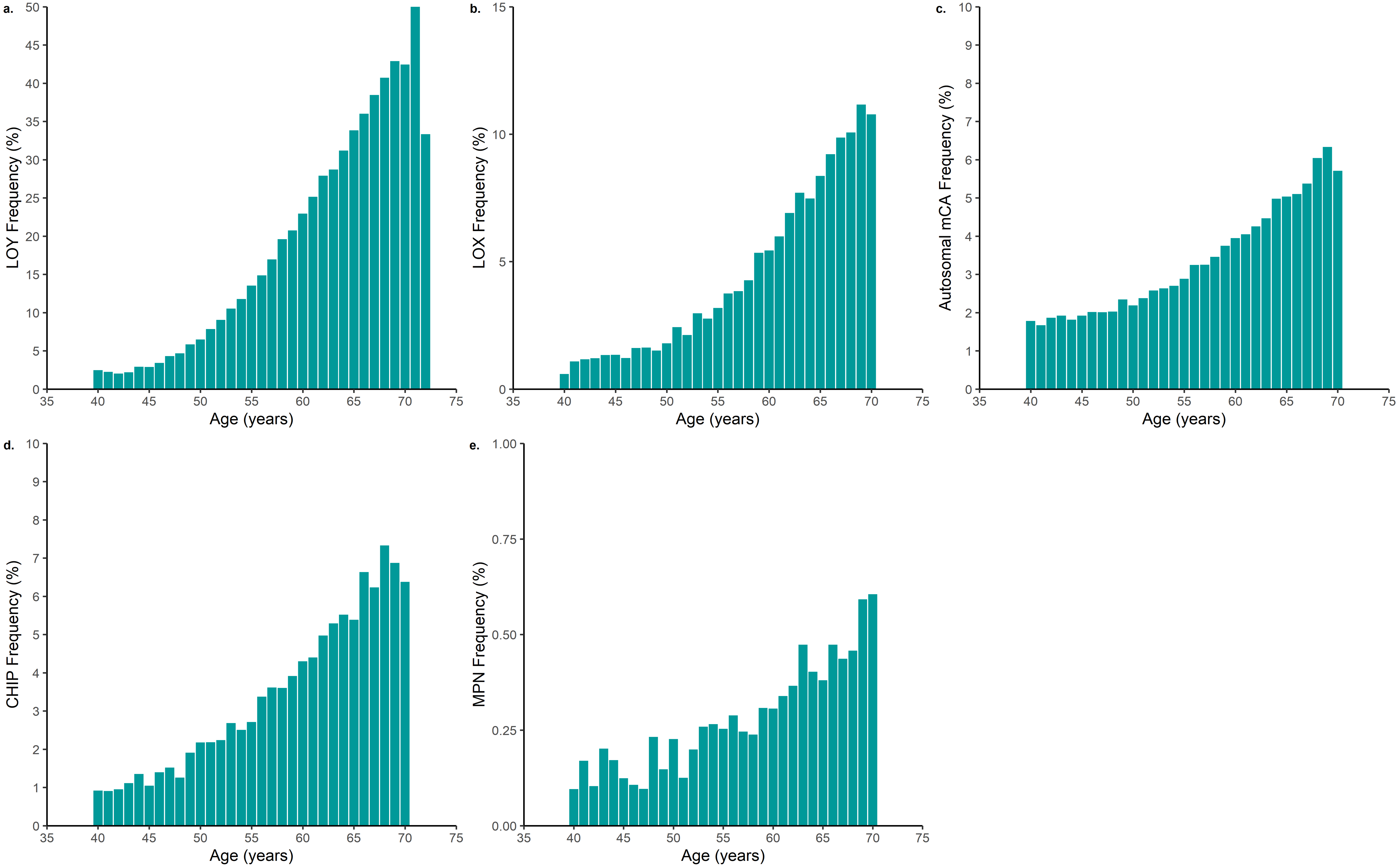


**Supplemental Figure 2**. Age distribution of each type of CH in the UK Biobank.

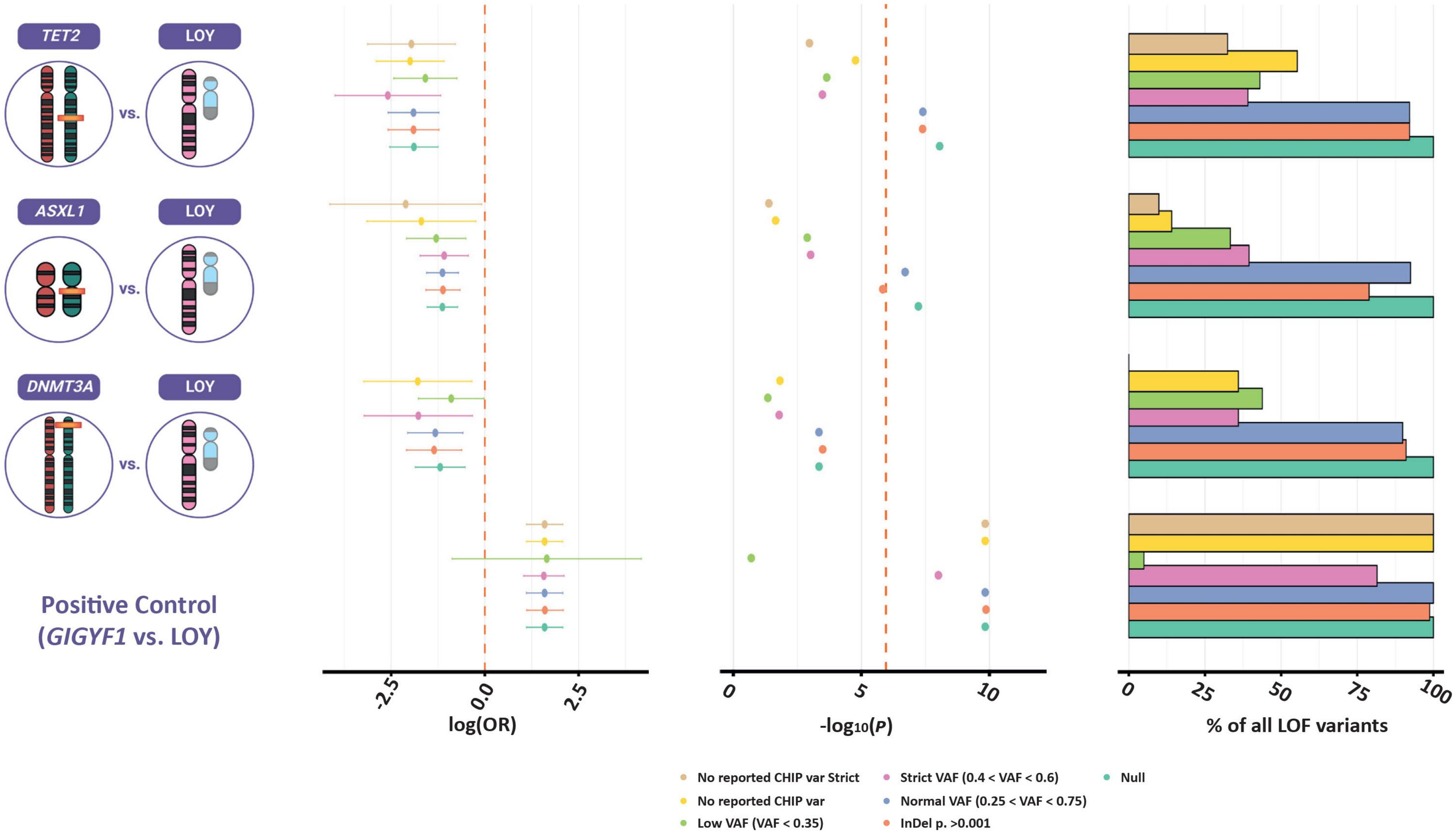


**Supplemental Figure 3**. Associations between CHIP loss of function variant carriers and LOY in three known CHIP genes, *TET2* (n = 193 carriers), *ASXL1* (n = 213 carriers), and *DNMT3A* (n = 89 carriers), and one control gene, *GIGYF1* (n = 81 carriers). Detailed results are given in **Supplemental Table 7**.

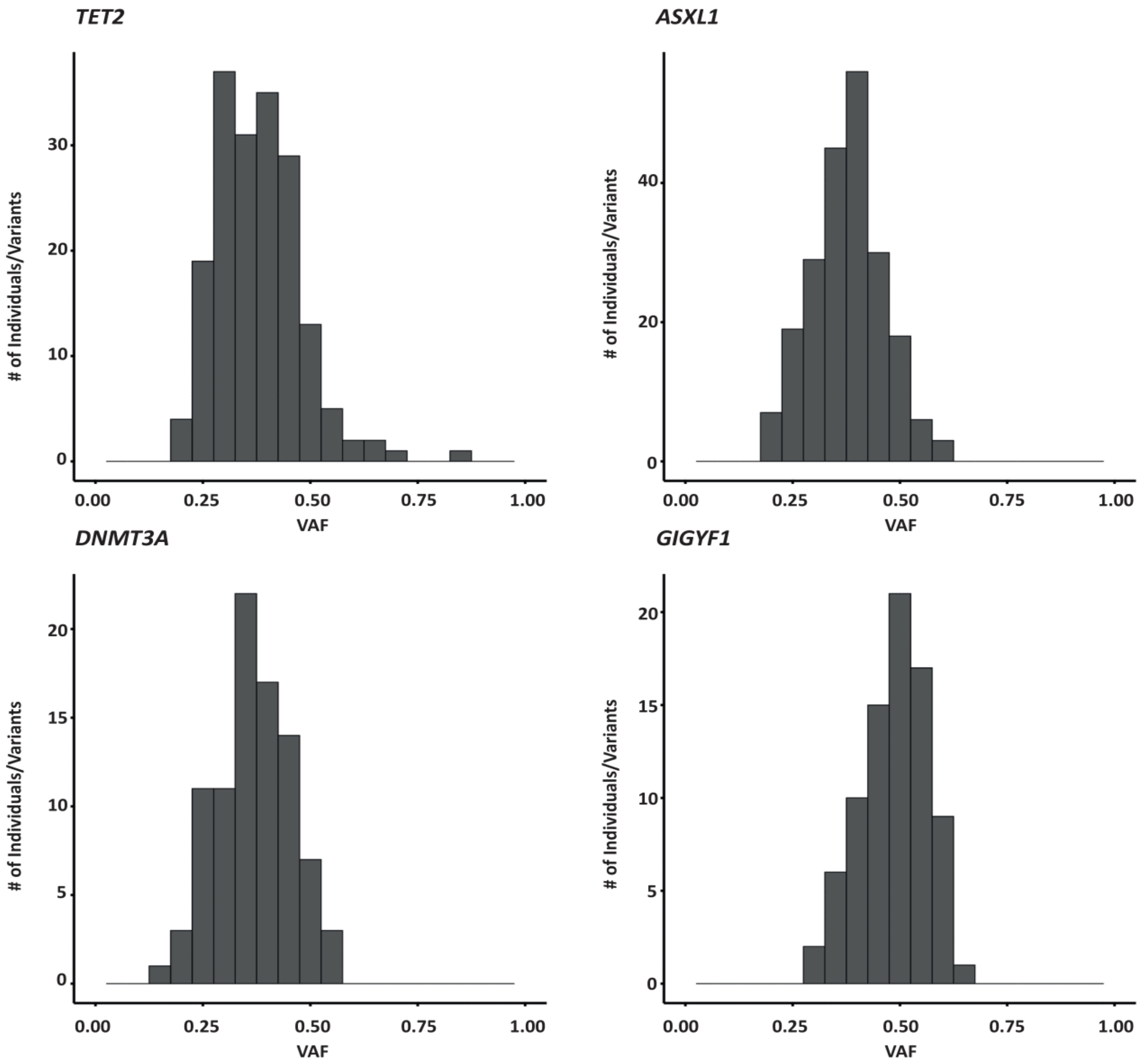


**Supplemental Figure 4**. Variant allele fraction (VAF) from the number of reads supporting the alternate and reference alleles for all carriers of protein truncating variants (PTV) at MAF < 0.1% in three known CHIP genes, *ASXL1* (n = 213 carriers), *DNMT3A* (n = 89 carriers), and *TET2* (n = 193 carriers), and one control gene, *GIGYF1* (n = 81 carriers).

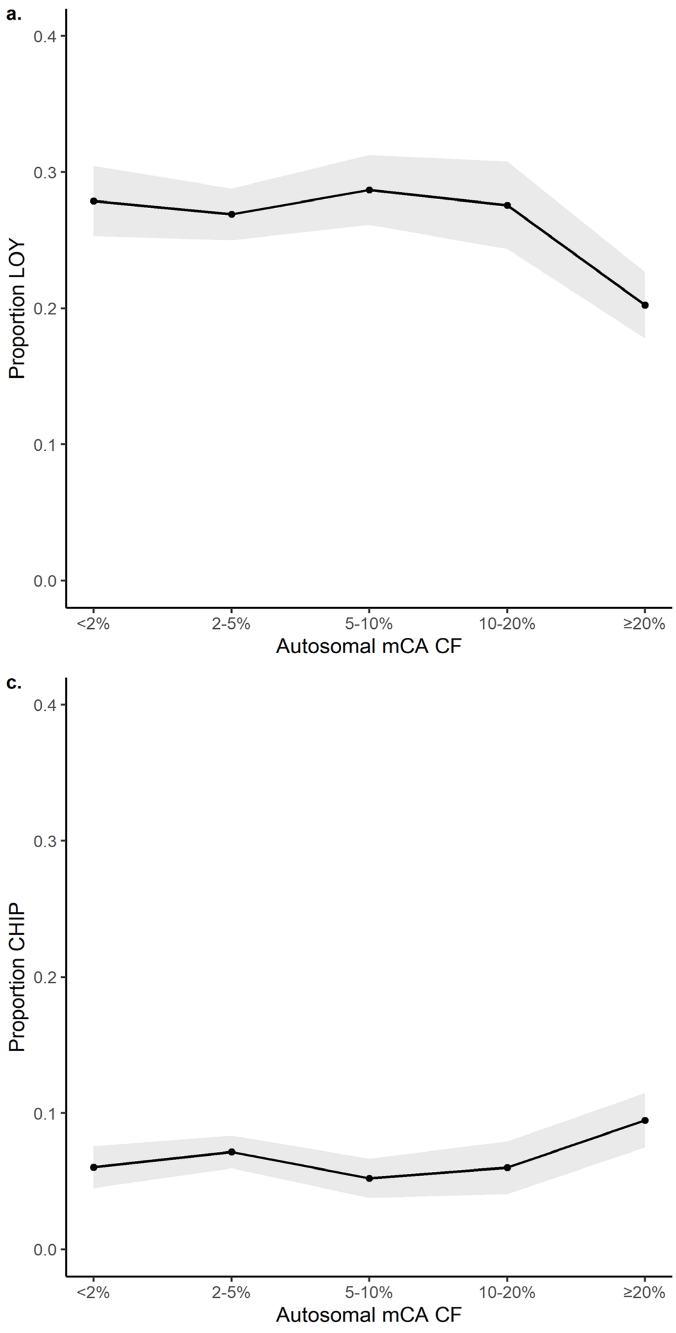


**Supplemental Figure 5**. Proportion of individuals with each type of CH by autosomal mCA cellular fraction. 95% confidence intervals (gray fill) are also plotted. Detailed plot data is given in **Supplemental Table 8**.

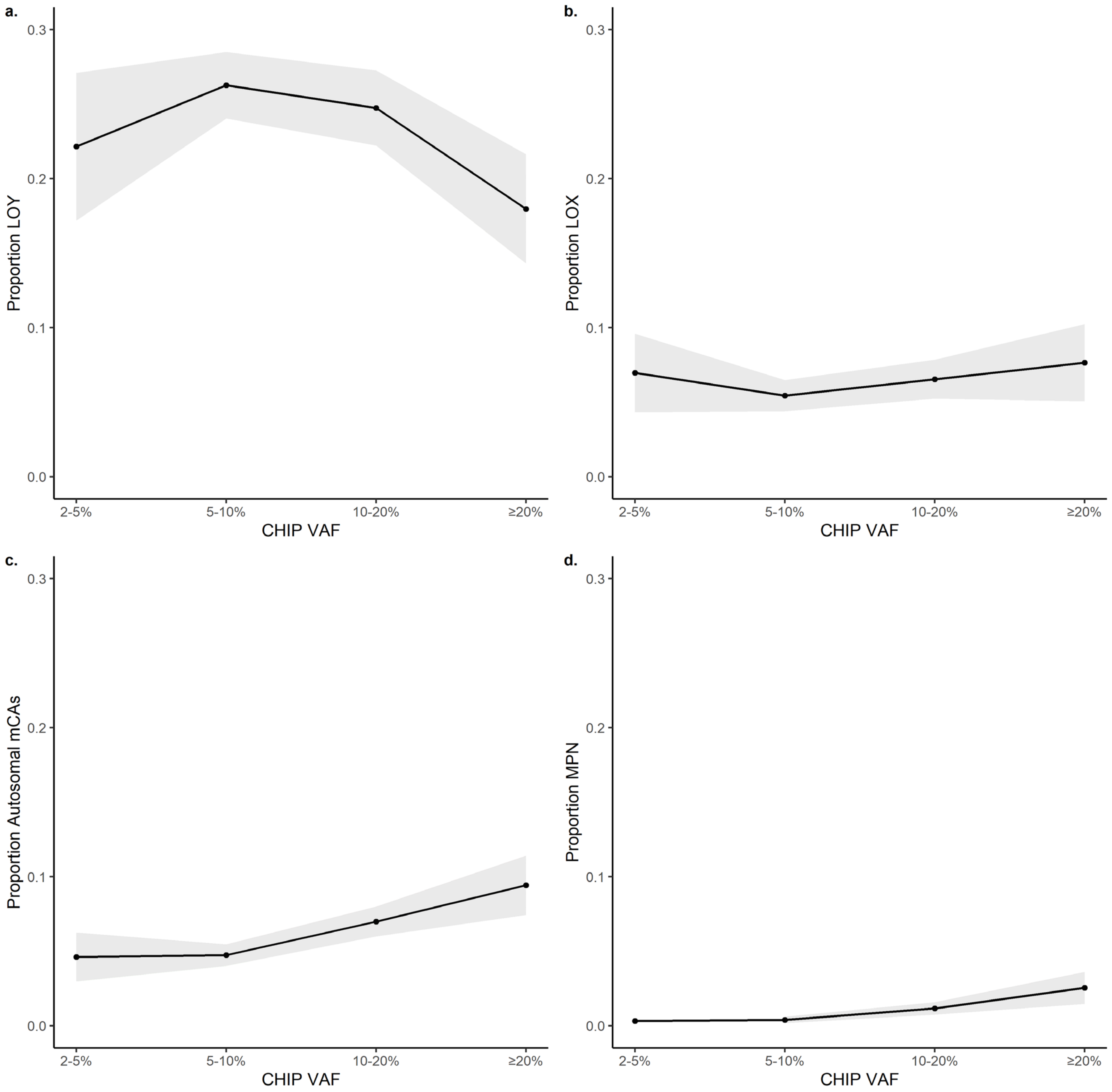


**Supplemental Figure 6**. Proportion of individuals with each type of CH by CHIP VAF. 95% confidence intervals (gray fill) are also plotted. Detailed plot data is given in **Supplemental Table 9**.

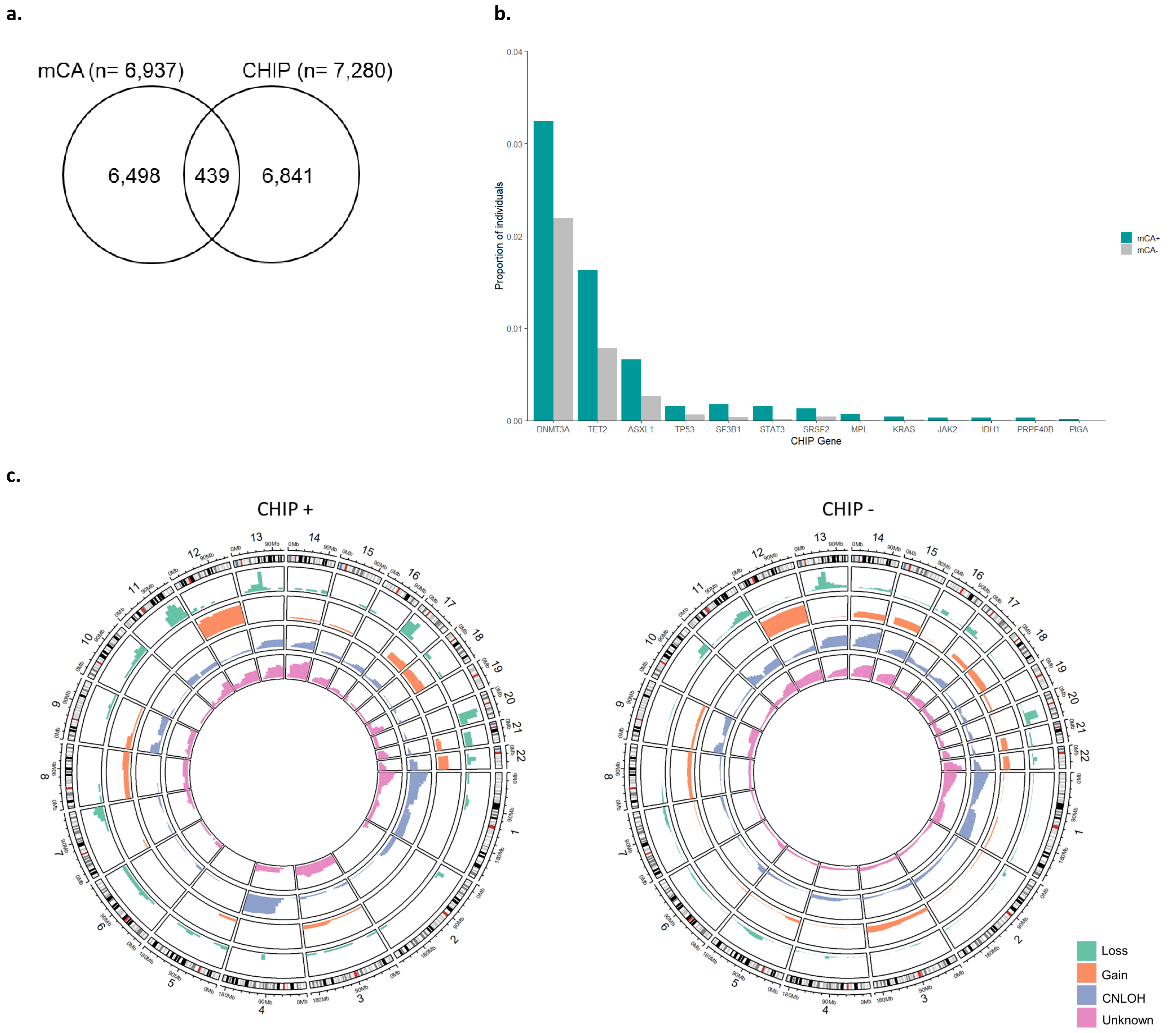


**Supplemental Figure 7.** (**a**) Overlap between autosomal mCAs and CHIP. The number of subjects from UK Biobank are given. (**b**) Proportion of CHIP gene mutations significantly enriched in subjects with autosomal mCAs. Frequencies for each CHIP mutation are given in **Supplemental Table 10**. (**c**) Circos plots of autosomal mCAs by CHIP mutation status. Frequency of autosomal mCAs by chromosome are given in **Supplemental Table 11**.

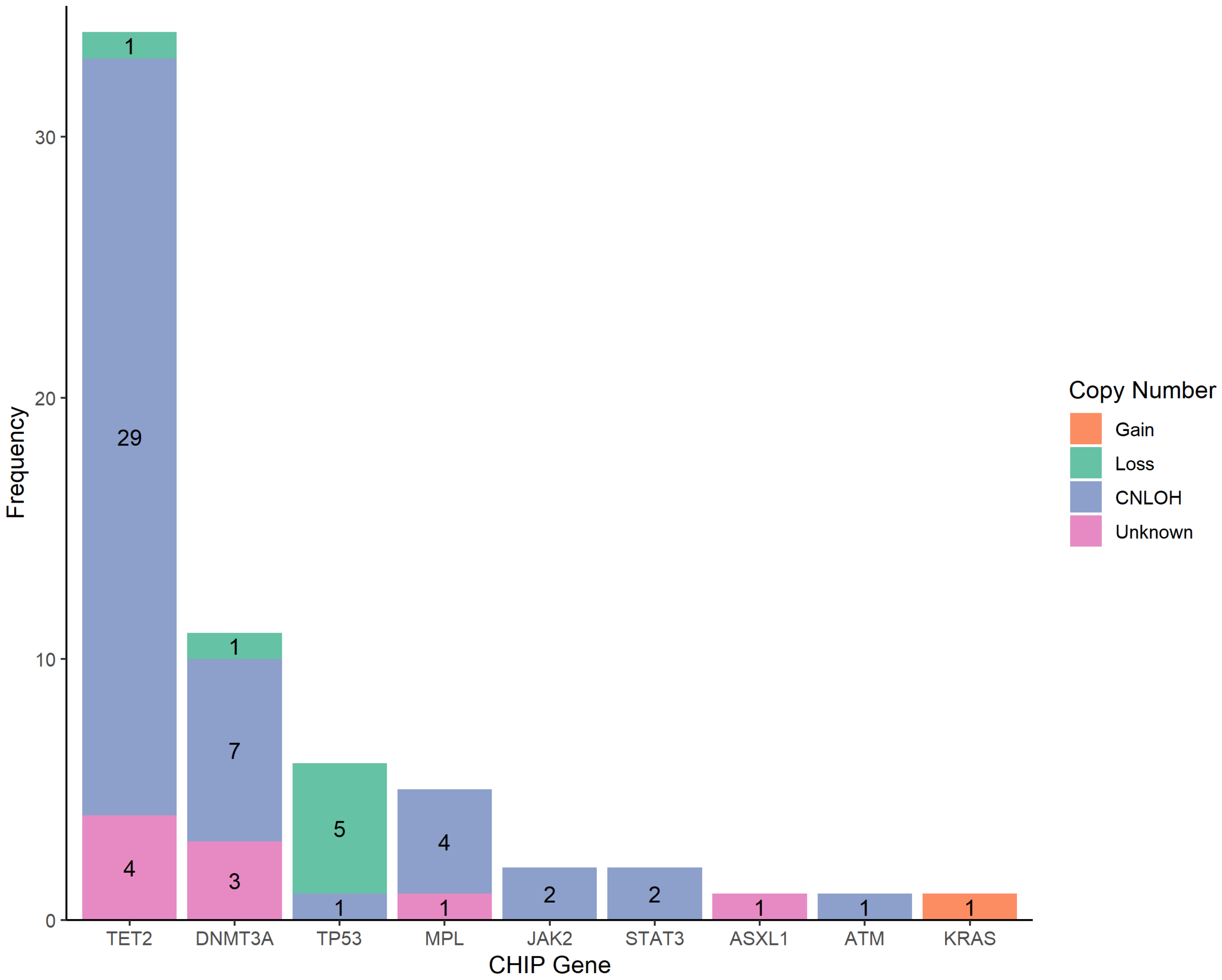


**Supplemental Figure 8**. Frequency of overlapping CHIP mutations and autosomal mCA events. 63 overlapping mutations were observed within 53 individuals. The y-axis depicts the frequency of each CHIP mutation. The frequency of each autosomal mCA copy number is labeled in each bar.

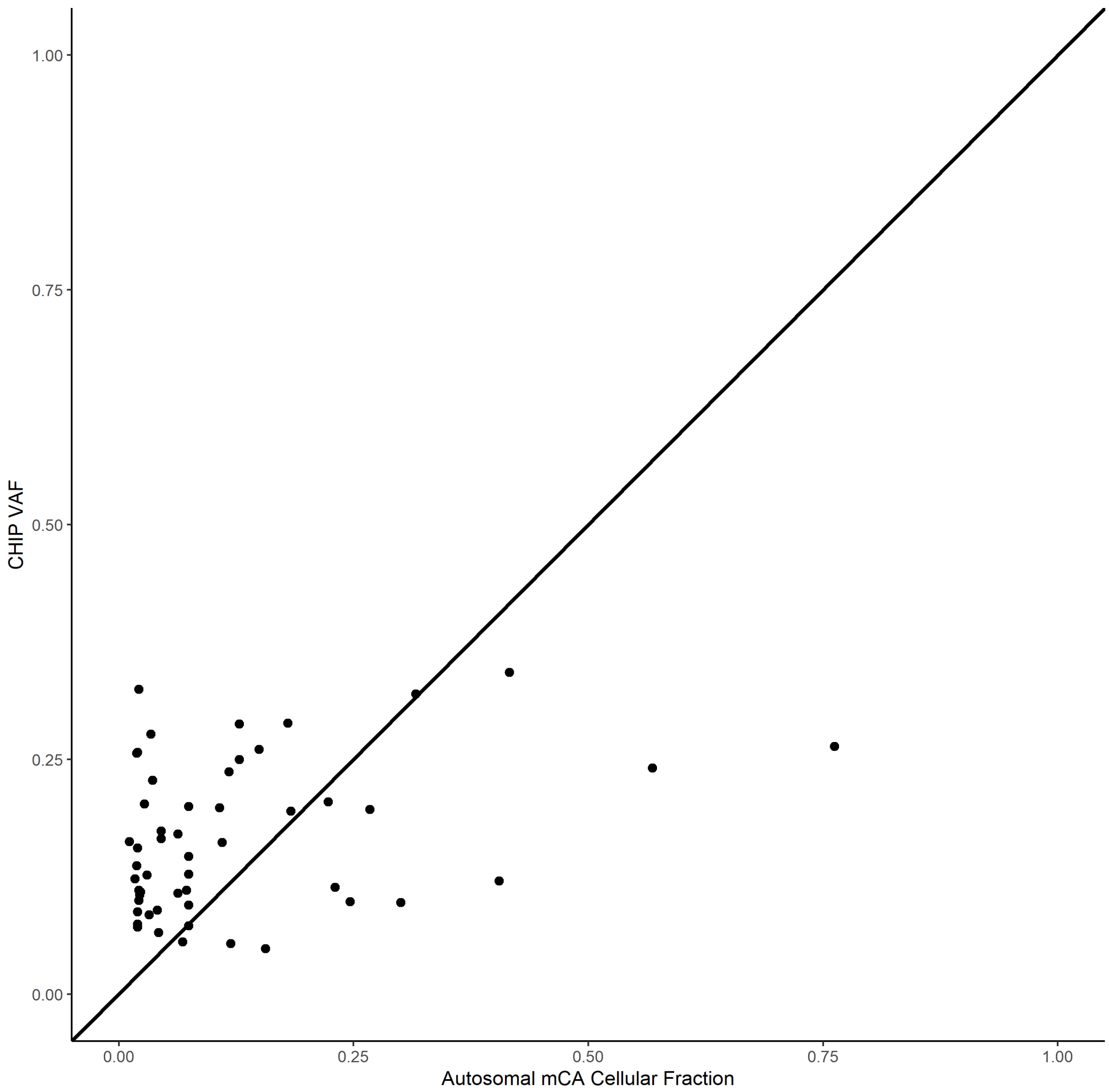


**Supplemental Figure 9**. Clonal evolution of CHIP and autosomal mCA mutations. The y-axis depicts the estimated CHIP VAF. The x-axis depicts estimated autosomal mCA cellular fraction. Autosomal mCA cellular fraction was missing for 9 events, due to unknown copy number state. CHIP VAF was larger than mCA cellular fraction for 41 (75.9%) overlapping mutations.

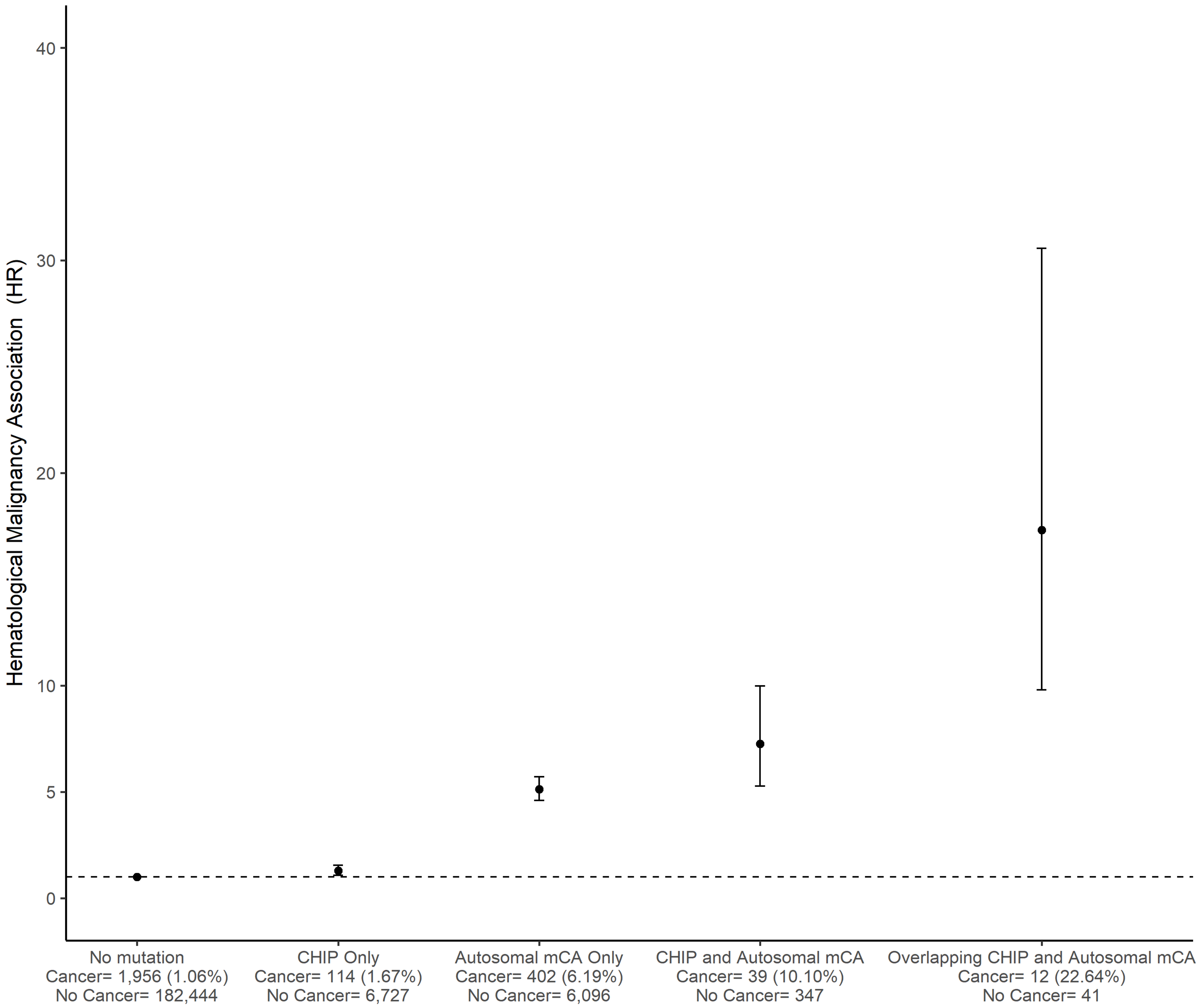


**Supplemental Figure 10**. Incident hematological malignancy association (HR) with 95% confidence intervals by CHIP and autosomal mCA status. The number of subjects with and without cancer within each group from UK Biobank are given. Associations are adjusted for age, age-squared, sex, and detailed smoking status. Associations are given in **Supplemental Table 12**.

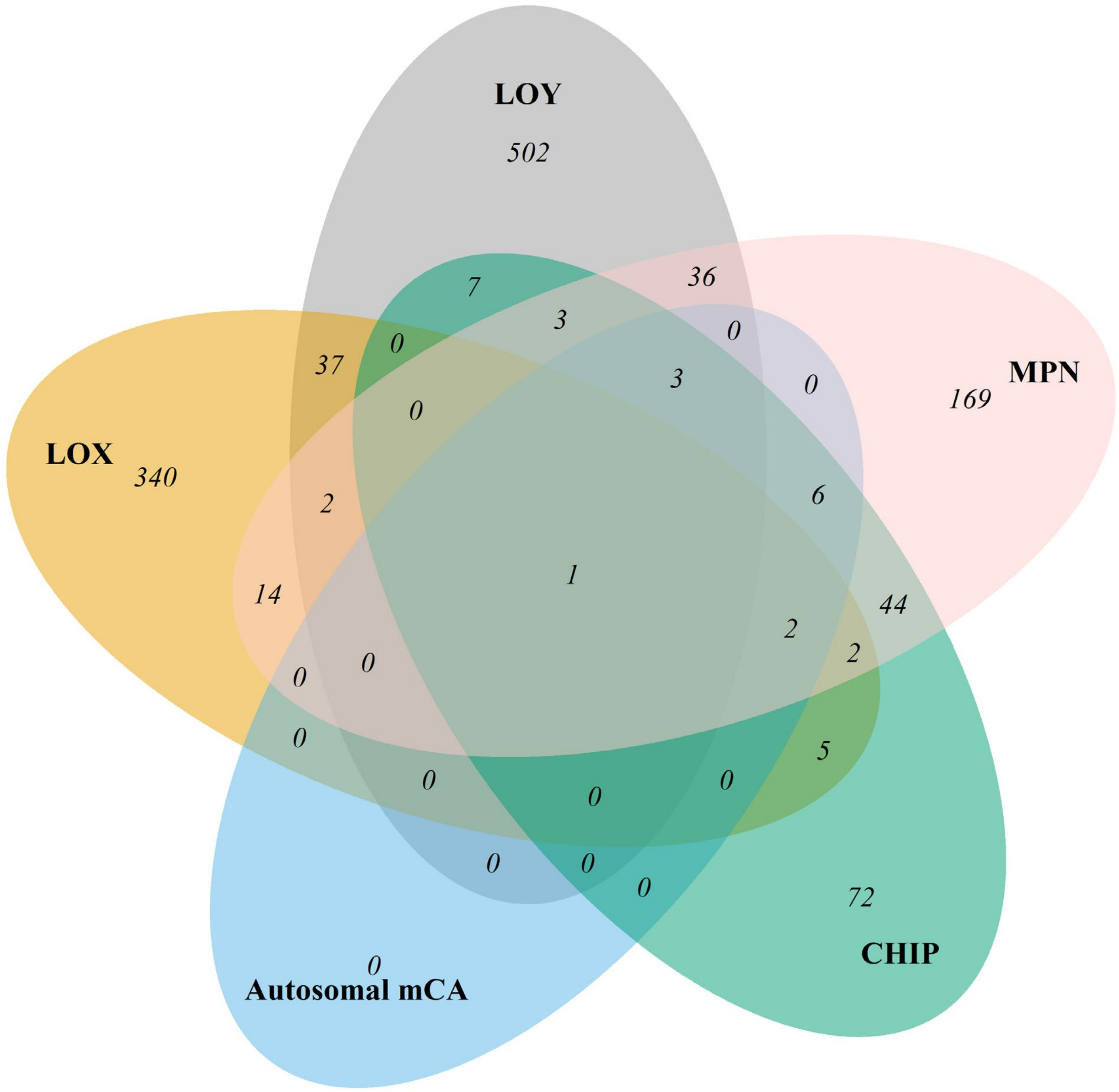


**Supplemental Figure 11**. Overlap of significant pathways and gene sets by type of CH. Detailed results are given in **Supplemental Tables 13-17**.

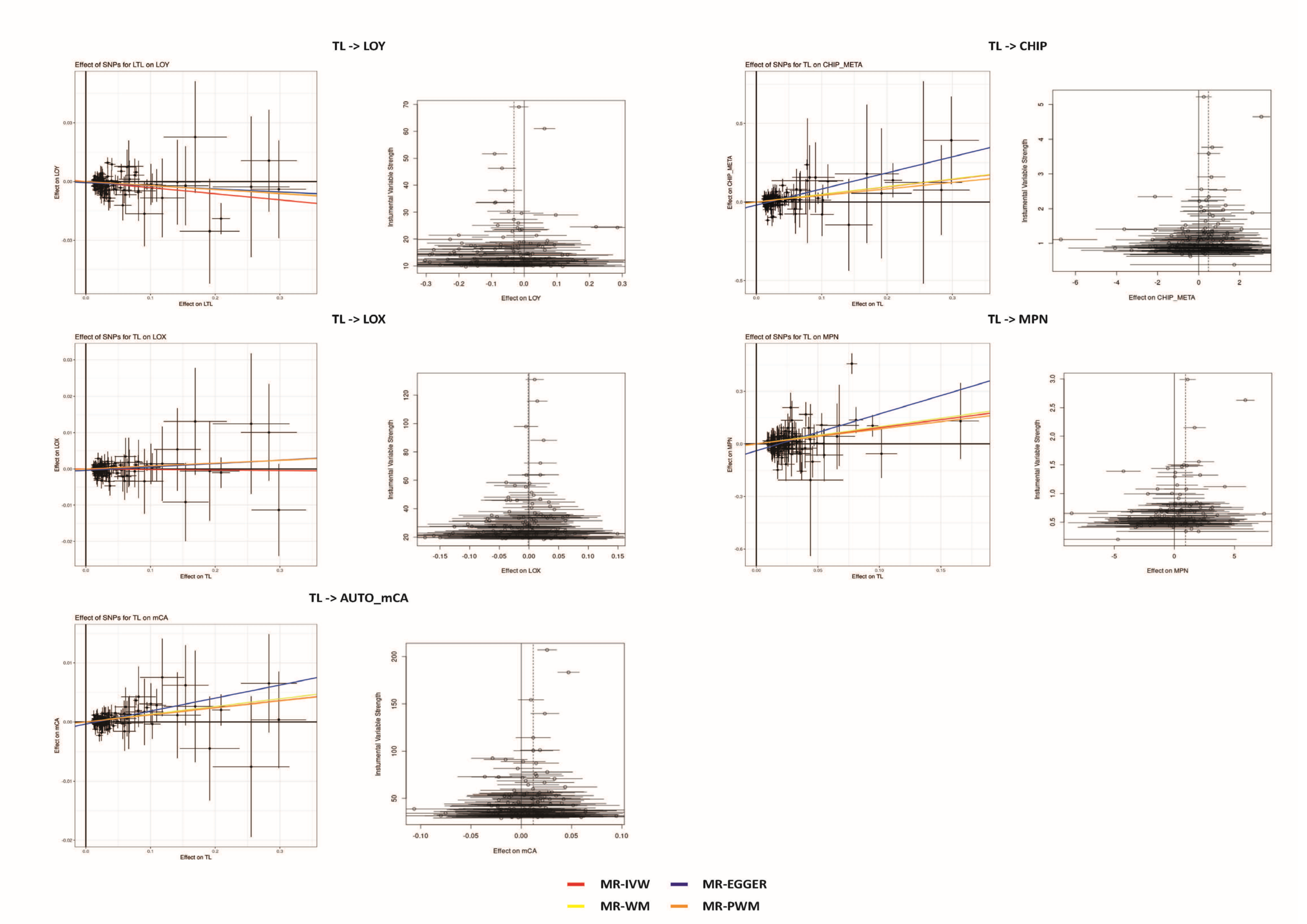


**Supplemental Figure 12**. Funnel and dosage plots for Mendelian randomization (MR) analyses using the TL instrument variable and each type of CH.

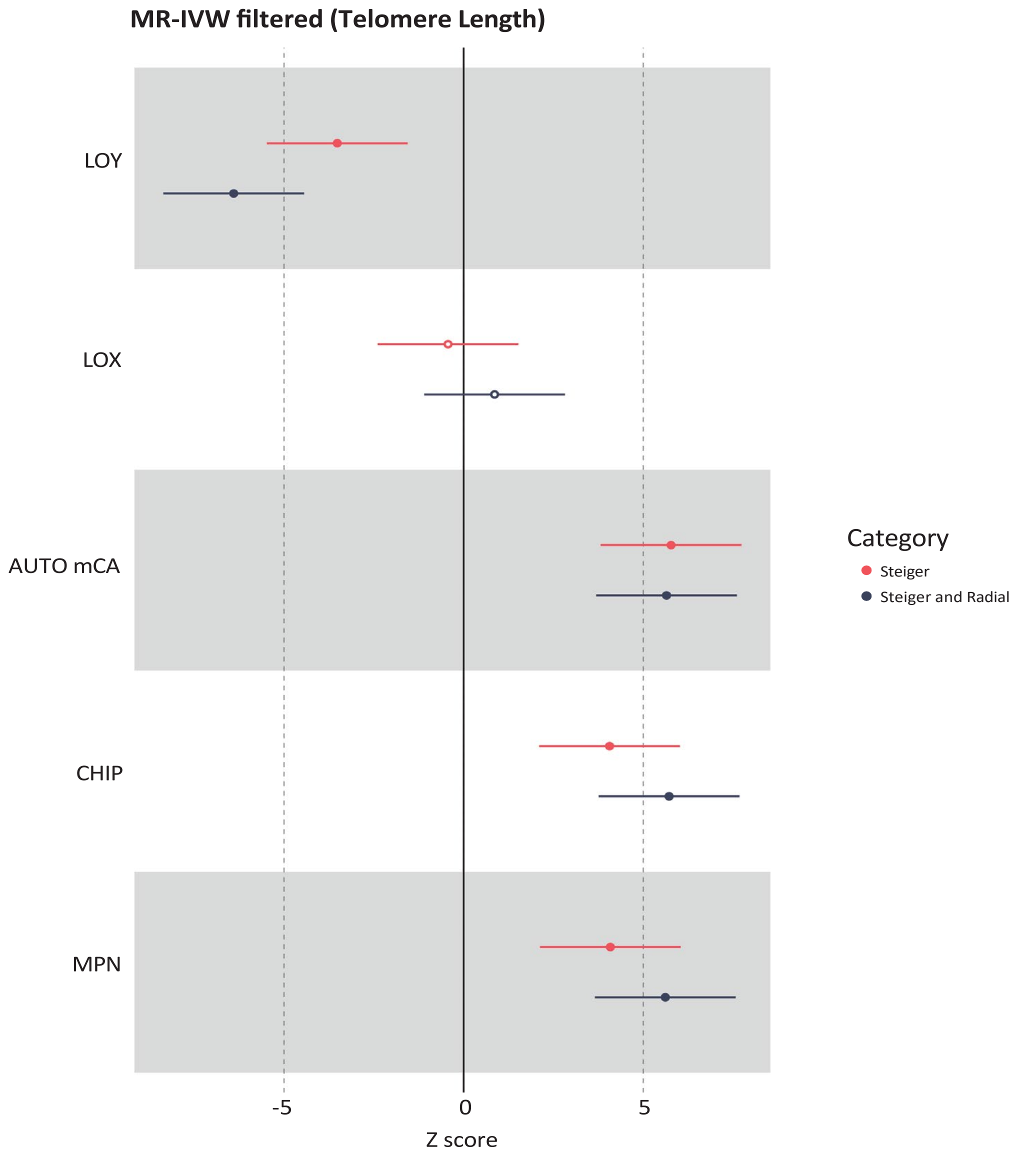


**Supplemental Figure 13**. MR inverse-variance weighted results between telomere length and each type of CH using telomere length instrumental SNPs with Steiger and Radial filters. Detailed results are given in **Supplemental Table 18**.

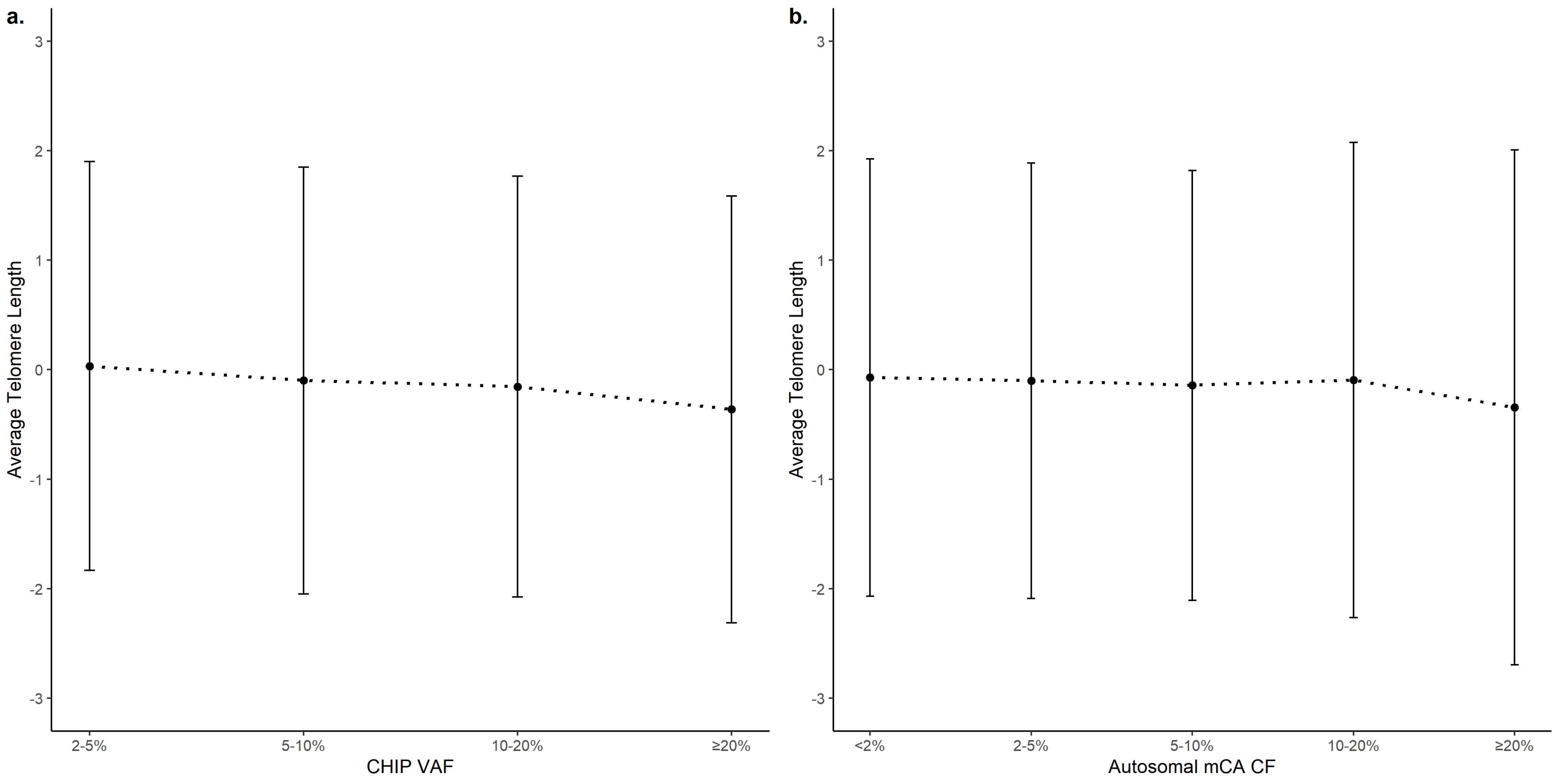


**Supplemental Figure 14**. (**a**) Average telomere length and 95% confidence intervals by CHIP VAF. Detailed results are given in **Supplemental Table 19**. (**b**) Average telomere length and 95% confidence intervals by autosomal mCA cellular fraction. Detailed results are given in **Supplemental Table 20**.

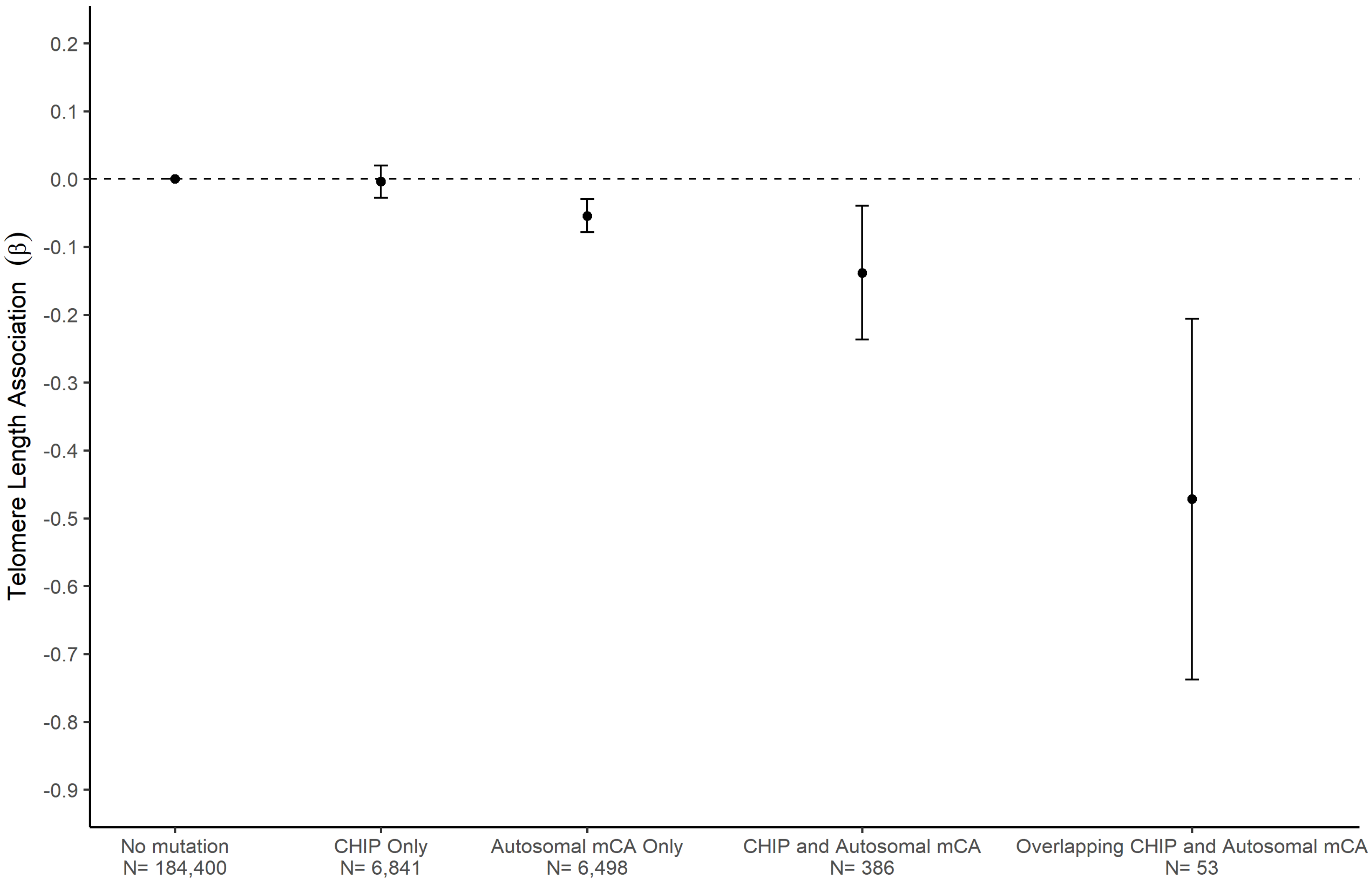


**Supplemental Figure 15**. Telomere length association (β) with 95% confidence intervals by CHIP and autosomal mCA status. The number of subjects within each group from UK Biobank are given. Associations are adjusted for age, age-squared, sex, and detailed smoking status. Detailed results are given in **Supplemental Table 21**.

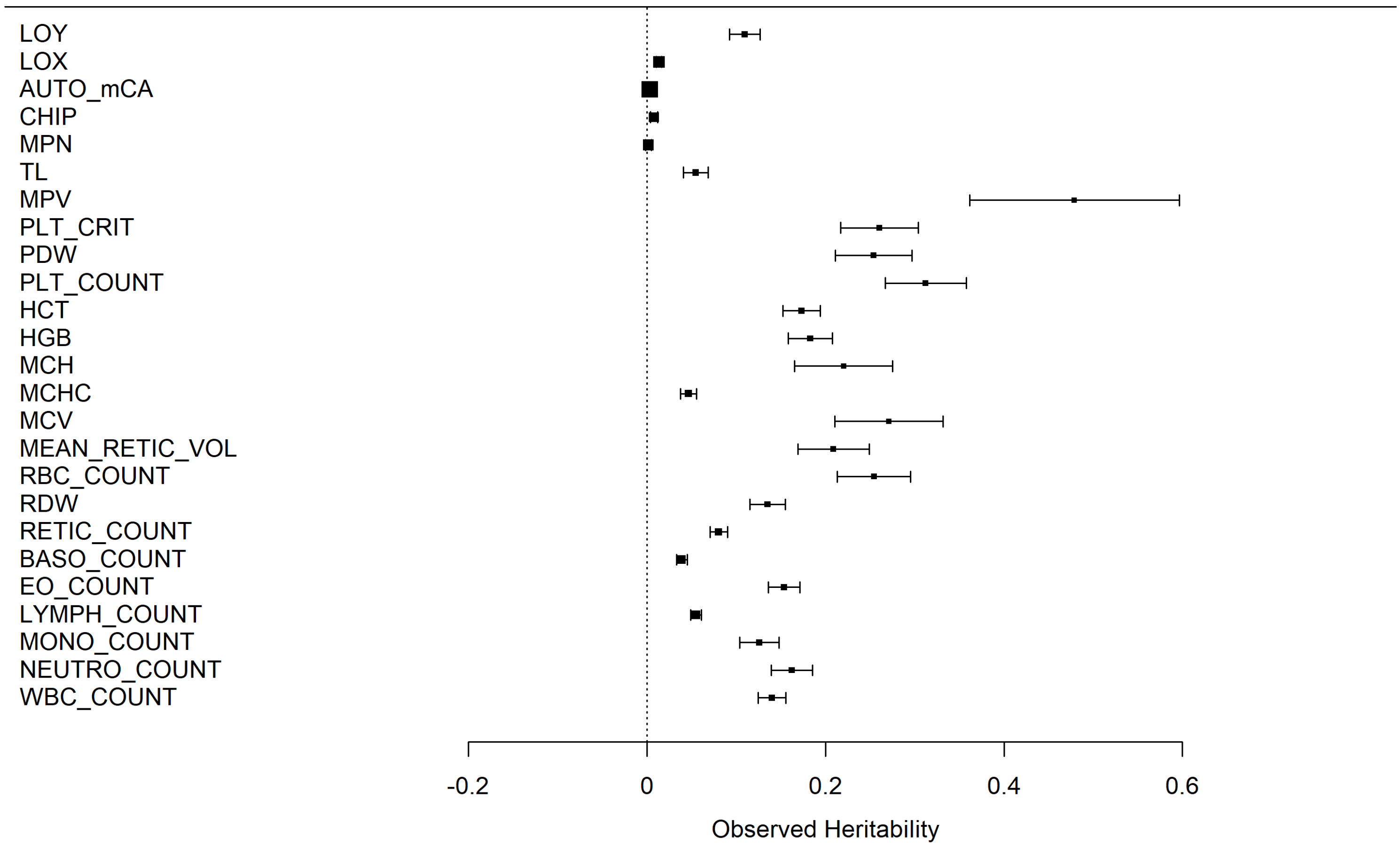


**Supplemental Figure 16**. Observed heritability for each type of CH, telomere length, and 19 blood cell traits. Detailed values are given in **Supplemental Table 28**.
